# CAF-1 deposits newly synthesized histones during DNA replication using distinct mechanisms on the leading and lagging strands

**DOI:** 10.1101/2022.10.14.512229

**Authors:** Clément Rouillon, Bruna V. Eckhardt, Leonie Kollenstart, Fabian Gruss, Alexander E.E. Verkennis, Inge Rondeel, Peter H.L. Krijger, Giulia Ricci, Alva Biran, Theo van Laar, Charlotte M. Delvaux de Fenffe, Georgiana Luppens, Pascal Albanese, Richard A. Scheltema, Wouter de Laat, Nynke H. Dekker, Anja Groth, Francesca Mattiroli

## Abstract

During every cell cycle, both the genome and the associated chromatin must be accurately replicated. Chromatin Assembly Factor-1 (CAF-1) is a key regulator of chromatin replication, but how CAF-1 cooperates with the DNA replication machinery is unknown. Here, we reveal that this crosstalk differs between the leading and lagging strand at replication forks. Using biochemical reconstitution, we show that DNA and histones promote CAF-1 recruitment to its binding partner PCNA and reveal that two CAF-1 complexes are required for efficient nucleosome assembly under these conditions. Remarkably, in the context of the replisome, CAF-1 competes with the leading strand DNA polymerase epsilon (Polε) for PCNA binding, but not with the lagging strand DNA polymerase Delta (Polδ). Yet, in cells, CAF-1 deposits newly synthesized histones equally on both daughter strands. Thus, on the leading strand, chromatin assembly by CAF-1 cannot occur simultaneously to DNA synthesis, while on the lagging strand both processes are coupled. We propose that these differences may facilitate distinct parental histone recycling mechanisms and accommodate the inherent asymmetry of DNA replication.

## INTRODUCTION

During every cell cycle, a new copy of the genome is made. At the same time, genomic chromatin organization must be replicated to ensure faithful transmission of the parental epigenetic state to both daughter cells after cell division. Therefore, genome and chromatin replication are tightly coupled and regulated by the concerted action of several dozens of proteins. Errors in both processes affect cell function; they can derail developmental programs or cause diseases, such as cancer (1–4).

DNA is replicated by the replisome, which is comprised of a core Cdc45-MCM-GINS (CMG) helicase complex, DNA polymerases and regulatory factors (5–9). Two distinct DNA polymerases function on the two daughter strands: DNA polymerase epsilon (Polε) acts on the leading strand, whereas DNA polymerase delta (Polδ) acts on the lagging strand (10–14). Because both DNA polymerases synthesize DNA in the 5’-3’ direction, the two strands are replicated via distinct mechanisms. Polε tightly binds the CMG and continuously extends the leading strand, while Polδ discontinuously synthesizes short Okazaki fragments, which are later processed and ligated on the lagging strand (13–17). Despite their mechanistic differences, both DNA polymerases require the processivity factor <u>P</u>roliferating <u>C</u>ell <u>N</u>uclear <u>A</u>ntigen (PCNA) for their function. PCNA is an essential homotrimeric clamp that encircles newly synthesized double-stranded DNA, tethering the DNA polymerases to DNA. It is abundant at replication forks where, in addition to the DNA polymerases, it binds many other factors involved in genome replication, chromatin assembly and the response to stress and damage (18–20).

Chromatin replication requires proteins that function as histone chaperones, which include replisome components with histone binding properties (i.e. MCM2, Polε, Polα and RPA) and *bona fide* histone chaperones that are recruited to the replisome (i.e. FACT, CAF-1 and ASF1) (21). These proteins coordinate the recycling of parental histones to spatially maintain the landscape of histone post-translational modifications. They also promote the incorporation of newly synthesized histones to preserve nucleosome density on the daughter DNA strands (2, 4). Replicated DNA is readily assembled into chromatin (22, 23), a process that constitutes the first critical step to the re-establishment of epigenetic modifications on histones genome-wide (2, 4, 24–28).

Chromatin Assembly Factor-1 (CAF-1) is a key regulator of chromatin assembly during DNA replication (29). CAF-1 deletion is lethal during vertebrate development (30–32), and transient CAF-1 depletion affects cell cycle progression and cell fate (27, 33–42). CAF-1 forms a heterotrimeric complex consisting of Cac1, Cac2 and Cac3 in yeast and p150, p60 and p48 in mammals. The complex chaperones newly synthesized histones H3-H4 and deposits them onto DNA at sites of DNA synthesis (43–48). CAF-1 activity at replication forks depends on its interaction with PCNA, which occurs via canonical <u>P</u>CNA <u>I</u>nteracting <u>P</u>eptides (PIPs) present on the large CAF-1 subunit (49–53). While the function of CAF-1 has been studied in cells and in the SV40 systems, a detailed bottom-up biochemical reconstitution to address the molecular mechanism by which CAF-1 assembles chromatin during DNA replication and its interplays with the replisome is still lacking.

Here we developed biochemical systems to study the crosstalk between CAF-1 and key components of the DNA replication machinery, combining our previous CAF-1 histone chaperone assays (54, 55) with primer extension assays and the recent *in vitro* reconstitutions of the eukaryotic replisome (8, 9). We find that CAF-1 recruitment to PCNA requires DNA and is modulated by histones. Two CAF-1 complexes bind PCNA and are necessary for PCNA-dependent nucleosome assembly. CAF-1 interaction with PCNA inhibits the activity of the leading-strand DNA polymerase Polε, but not of the lagging-strand polymerase Polδ. Yet, in cells, we show that CAF-1 deposits histones equally on the leading and lagging strands during DNA replication. Thus, our work reveals an unexpected difference in the crosstalk between CAF-1, PCNA and the two replicative polymerases, indicating different mechanisms for the coupling of nucleosome assembly to DNA synthesis on the two daughter strands.

## RESULTS

### CAF-1 recruitment to PCNA requires DNA

We first set out to study the interaction between CAF-1 and PCNA in the context of DNA, as this is the context in which the CAF-1-PCNA interaction occur during DNA replication. Therefore, we loaded PCNA onto nicked plasmids using the ATP-dependent clamp loader RFC1-5 (70), and separated DNA-loaded from free PCNA on a size exclusion column (SEC) (Supplementary Figure S1A) (71). When adding CAF-1, we observed that the three CAF-1 subunits co-eluted with DNA-loaded PCNA, suggesting the formation of a CAF-1-PCNA-plasmid complex (Figure 1A and Supplementary Figure S1B-C). As CAF-1 uses PIPs to bind PCNA in cells (39, 51–53), we introduced mutations in these domains to test their importance in our *in vitro* system (Supplementary Figure S1D). The mutant CAF-1_PIP** no longer bound to DNA-loaded PCNA (Figure 1B and Supplementary Figure S1C), confirming that our *in vitro* reconstitution recapitulates the physiological determinants of the CAF-1-PCNA interaction.

**Figure 1:**
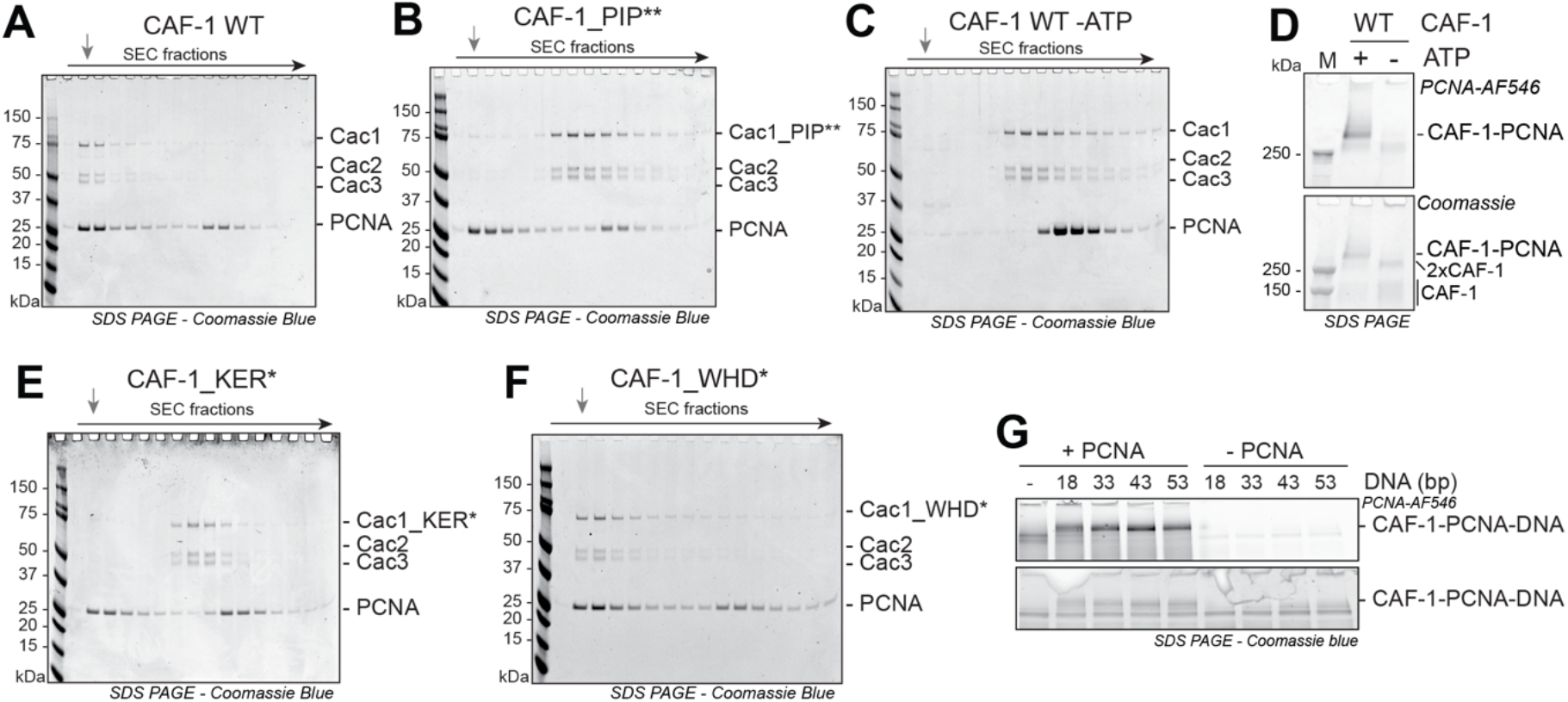
DNA and histones control CAF-1 recruitment to DNA-loaded PCNA. **A-C)** SDS PAGE following separation on SEC of a CAF-1-PCNA binding reaction on DNA plasmids using WT CAF-1 (A), a CAF-1_PIP** mutant (B), or WT CAF-1 in absence of ATP (C). The grey arrow indicates the elution volume of the plasmid DNA. Chromatograms are shown in Supplementary Figure S1C. **D)** SDS PAGE of crosslinking reactions of fluorescent PCNA (3 μM) and CAF-1 (1.5 μM), RFC (150 nM) and nicked pUC19 plasmid (300 nM) after nuclease digestion. **E-F)** Coomassie-stained SDS PAGE following SEC of a CAF-1-PCNA binding reaction on DNA plasmids using CAF-1_KER* (E) and CAF-1_WHD* (F) mutants. **G)** Crosslinking experiment between CAF-1 (3 μM) and labeled PCNA (4.5 μM) on DNA fragments (1.5 μM) of various sizes. RFC and ATP were not added to actively load PCNA and DNA was not digested in these reactions. Full gels are shown in Supplementary Figure S1I.

Next, we investigated how DNA contributes to the CAF-1-PCNA interaction. CAF-1 did not co-elute with PCNA in the absence of DNA (Supplementary Figure S1E) or when PCNA was not loaded onto DNA (i.e. by omission of ATP) (Figure 1C and Supplementary Figure S1C), suggesting that DNA is required for the CAF-1-PCNA interaction. To confirm that the interaction between CAF-1 and PCNA is DNA-dependent, we crosslinked CAF-1 to fluorescently labeled PCNA on nicked DNA plasmids using glutaraldehyde in the presence or absence of ATP, followed by nuclease digestion and SDS-PAGE analysis to determine if more transient protein-protein complexes are formed in solution, which may be lost during the SEC purification. Again, we observed no significant CAF-1-PCNA complexes when PCNA was not loaded onto DNA (Figure 1D). These results indicate that DNA is required for a stable interaction between CAF-1 and PCNA.

CAF-1 contains two DNA binding regions in its large Cac1 subunit: the Lys-Glu-Arg rich (or KER) region located at the N-terminus, which is flanked by the PIPs, and the winged-helix domain (WHD) at the C-terminus (Supplementary Figure S1D). Either domain is required for CAF-1 function in cells (48, 53, 72, 73), but their relative role in CAF-1 mechanism remains unclear, as both domains must be mutated simultaneously in order to disrupt CAF-1 activity *in vitro* in the absence of PCNA (54). We thus tested whether these domains contributed to the DNA-dependent interaction of CAF-1 to PCNA. Deletion of the KER domain or its mutation into a neutral unstructured sequence (CAF-1_ΔKER and CAF-1_KER*, respectively) abrogated the interaction between CAF-1 and DNA-loaded PCNA, similarly to the effect of the CAF-1_PIP** mutant (Figure 1E and Supplementary Figure S1F). However, mutations in the WHD (CAF-1_WHD*) had no effect on binding to DNA-loaded PCNA (Figure 1F). These results indicate that the CAF-1 KER domain, but not the WHD, is critical for the formation of a stable CAF-1-PCNA complex on DNA.

Having established that DNA is required for the CAF-1-PCNA interaction, we investigated whether there is a minimum DNA length required to promote this interaction. We first confirm that CAF-1 binds 10-fold more weakly to a 18 bp DNA (Kd > 2 μM) than to a 33, 43 or 53 bp DNA (Kd = 0,33, 0,23 and 0,18 μM respectively) (Supplementary Figure S1G), in line with previous observations (74). Mutations in the KER domain strongly inhibit the CAF-1-DNA interaction, while WHD mutations have a minor effect (Supplementary Figure S1H). Complex formation was less efficient on the 18 bp DNA fragment, where PCNA can load in the absence of RFC, than on the longer 43 and 53 bp DNAs (Figure 1G and Supplementary Figure S1I). This suggests that a minimum of ±30 bp need to be exposed for CAF-1 to stably bind PCNA on DNA. Notably, the Alphafold model of the Cac1-KER domain (residues 128-226) predicts a long helical structure of ^~^145 Å, which corresponds to the length of ^~^44 bp of duplex DNA (Supplementary Figure S1J). This domain displays a positively charged surface along its helical arrangement, which may structurally explain the link we observe between DNA length and CAF-1 binding, assuming that this surface interacts with the negatively charged phosphate backbone of the DNA via electrostatic interactions. Overall, these observations suggest that the CAF-1-PCNA interaction on DNA is stabilized by DNA of at least ^~^30 bp via the KER domain in CAF-1.

### Reconstitution of PCNA-dependent nucleosome assembly by CAF-1

In cells, PCNA directs CAF-1-mediated chromatin assembly (43, 49–51). However, we and others have recently shown that CAF-1 is able to assemble nucleosomes *in vitro* in the absence of other factors (54, 74, 75). To determine how the presence of PCNA affects the histone chaperone activity of CAF-1, we set out to develop a nucleosome assembly assay that recapitulates the PCNA dependency of CAF-1 activity observed *in vivo*. The challenge is to differentiate between PCNA-dependent and PCNA-independent (i.e. purely DNA driven (54)) activity of CAF-1. To overcome this challenge, we mixed the nicked plasmid where PCNA can be loaded, with a competitor supercoiled plasmid where PCNA-independent reactions take place. Following a PCNA loading step, we added CAF-1-H3-H4 complexes to promote tetramer deposition followed by the addition of fluorescently labeled H2A-H2B, which associate with tetrasomes *in vitro* to form nucleosomes. Fluorescence-based readouts on native gels combined with micrococcal nuclease (MNase) digestion are used to measure nucleosome assembly on each of the plasmids. We named this setup PCNA-NAQ assay, based on our previously established <u>N</u>ucleosome <u>A</u>ssembly and <u>Q</u>uantitation (NAQ) assay (54, 62).

We first established that the PCNA-NAQ assay measures PCNA-dependent and -independent CAF-1 activity. Efficient nucleosome assembly (monitored by an increase in H2B fluorescence) on the nicked plasmid was observed only when PCNA is loaded on DNA and CAF-1 is present (Figure 2A). When PCNA loading was blocked by the omission of ATP, PCNA or RFC (Figure 2A), the histone fluorescence signal shifted to the supercoiled plasmid, confirming that the signal on the nicked plasmid is dependent on PCNA. As expected, omission of CAF-1 led to a drastic reduction of histone deposition (Figure 2A), reinforcing that CAF-1 is the nucleosome assembly machinery on both plasmids in our reconstitution. No nucleosomes were formed upon omission of either histones H3-H4 or H2A-H2B (Supplementary Figure S2A). Moreover, using labeled H3-H4 instead of H2A-H2B did not affect the results (Supplementary Figure S2B), confirming that our signal is a *bona fide* measure of assembled nucleosomes. Quantification of the histone fluorescence signal on the nicked plasmid compared to total histone signal showed that roughly 50% of the nucleosomes are assembled in a PCNA-dependent manner, when PCNA is loaded (Figure 2B). This is reduced to roughly 20% when PCNA was not loaded onto DNA (Figure 2B). These observations were confirmed using next generation sequencing approaches of the MNase products, when we used plasmids with distinct DNA sequences which allowed us to map relative nucleosome assembly and positioning (Supplementary Figure S2C-F). Thus, we developed a new method to study the PCNA-dependent nucleosome assembly function of CAF-1, where we can distinguish and quantify the PCNA-dependent or PCNA-independent activities of this histone chaperone complex.

**Figure 2:**
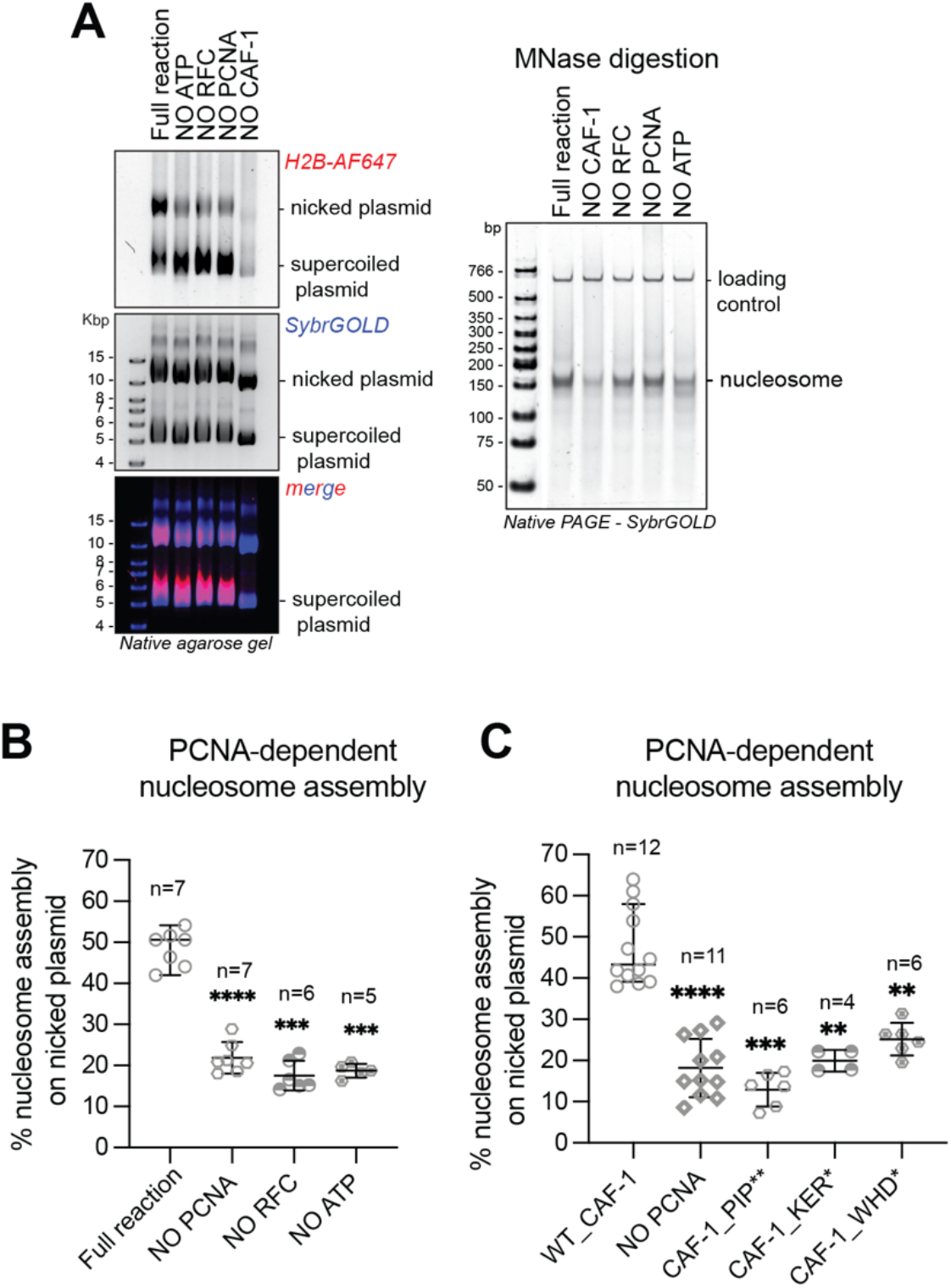
The WHD of CAF-1 controls PCNA-dependent nucleosome assembly. **A)** (Left) Native agarose gel of PCNA-NAQ assay reactions. A reaction containing all components, and reactions where we removed either ATP, RFC, PCNA or CAF-1 are shown. Fluorescence signal for H2B-T112C labeled with AF647 (H2B-AF647) or DNA (SybrGOLD), and their overlay are shown. H2B fluorescence on the nicked plasmid (top panel) represent PCNA-dependent histone deposition. (Right) Native PAGE stained with SybrGOLD to detect protected DNA fragments following MNase digestion of samples in A. 150bp DNA fragments are characteristic of nucleosomal DNA, a 621bp loading control is used to monitor DNA retrieval during the purification procedure. **B)** Quantification of the H2B fluorescence signal on the nicked plasmid relative to the total H2B signal in each lane in panel A as a measure of PCNA-dependent nucleosome assembly. **C)** Quantification of the PCNA-dependent nucleosome assembly activity for CAF-1_PIP**, CAF-1 KER* and CAF-1_WHD*. Means ±SD is shown, * p<0.05, ** p<0.01, *** p<0.001, **** p<0.0001 (one-way ANOVA comparing WT CAF-1 to control conditions (B) or each mutant (C)). Gels are shown in Supplementary Figure S2G.

We then used this method to understand how CAF-1 assembles nucleosomes when bound to PCNA. We first tested if mutations in the KER domain or PIPs of Cac1, which are important for recruitment to DNA-loaded PCNA (Figure 1B, 1E), affected its PCNA-mediated activity. As expected, CAF-1_KER* and CAF-1_PIP** showed a reduction specifically in PCNA-dependent nucleosome assembly (Figure 2C and Supplementary Figure S2G), while the overall activity of the mutant complexes was not affected as seen by the consistent level of MNase-protected nucleosome fragments (Supplementary Figure S2G). This confirms that CAF-1 recruitment is necessary for PCNA-dependent nucleosome formation in our PCNA-NAQ assay, further validating the role of these domains in the CAF-1-PCNA interaction. Strikingly, the CAF-1_WHD* mutant also showed a decrease in PCNA-dependent nucleosome assembly activity (Figure 2C and Supplementary Figure S2G), despite being able to bind DNA-loaded PCNA (Figure 1F) and being fully active in nucleosome assembly in absence of PCNA as shown by the MNase digestion products (Supplementary Figure S2G). This demonstrates that the WHD domain is important for PCNA-dependent CAF-1 activity specifically. Our observations explain why WHD mutations affect chromatin assembly during DNA replication in yeast cells (53, 54, 73), and why previous *in vitro* reconstitutions that omitted PCNA were unable to recapitulate loss of function of this mutant (54, 74). In summary, we show that the WHD domain in CAF-1 is important for the PCNA-dependent nucleosome assembly function of the complex.

### Two CAF-1 complexes bind PCNA to assemble nucleosomes

Two CAF-1 complexes are required to assemble one nucleosome in the absence of PCNA (54, 74). To understand how CAF-1 assembles nucleosomes when bound to PCNA, we therefore set out to study the stoichiometry of the CAF-1-PCNA complex on DNA. To this end, we used protein-protein crosslinking followed by nuclease digestion and SEC to analyze complexes in solution. These reactions elute with two peaks of equal distribution (Figure 3A). We collected fractions from these peaks and analyzed them by mass photometry to determine their composition (76). We found that Peak1 (Figure 3A) contained CAF-1-PCNA complexes corresponding to predominantly two CAF-1 per PCNA trimer (^~^430 kDa), and a lower amount of three CAF-1 per PCNA trimer (^~^590 kDa) (Figure 3B), while Peak2 contained mostly free unbound CAF-1 (^~^190 kDa) and a small fraction of complexes containing one CAF-1 per PCNA trimer (^~^285 kDa) (Figure 3B). These data indicate that the CAF-1-PCNA complex mainly assembles in a 2:1 (CAF-1 to PCNA trimer) stoichiometry on DNA, and to a lesser extent, can form 3:1 or 1:1 assemblies.

**Figure 3:**
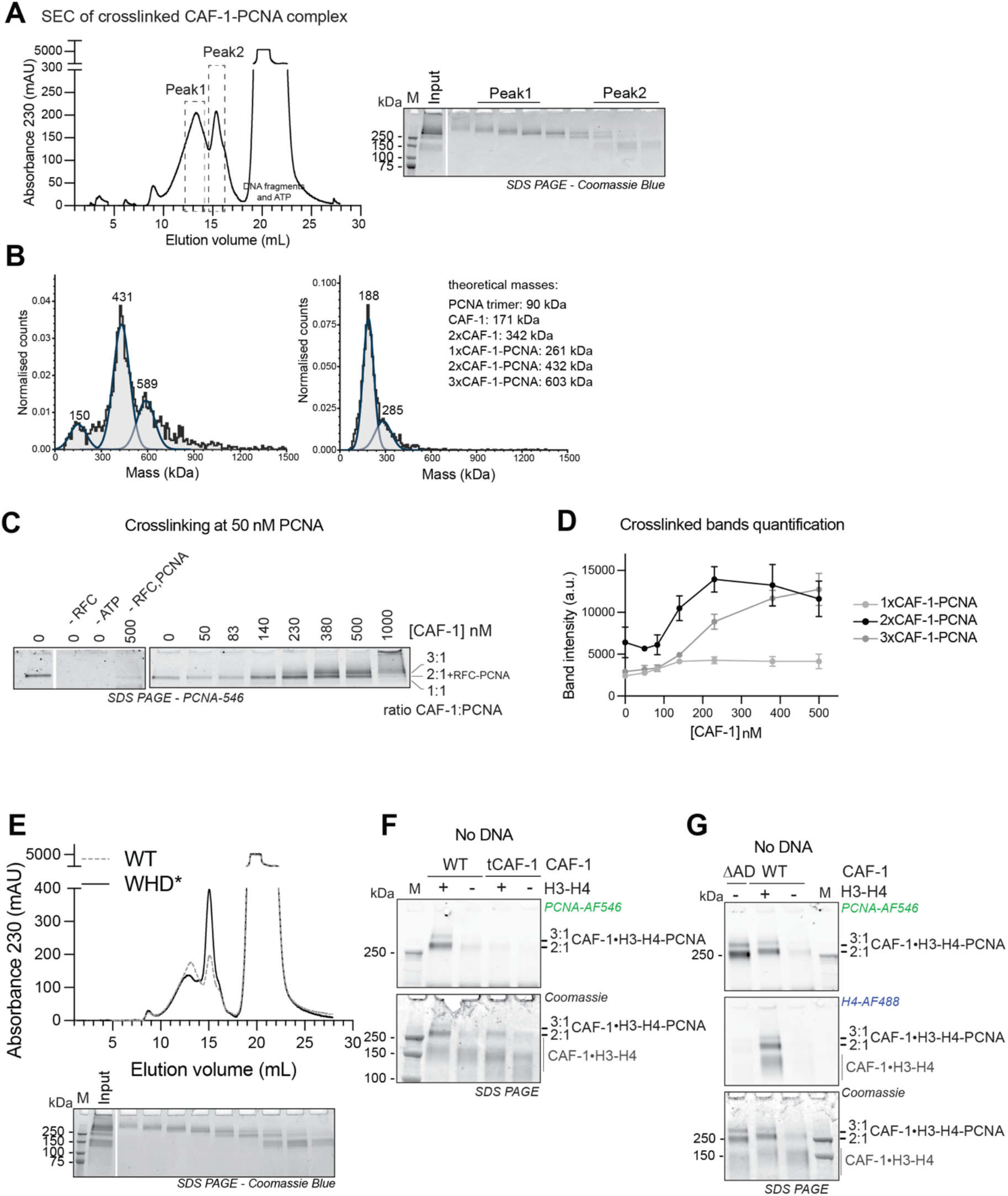
Two CAF-1 complexes bind to DNA-loaded PCNA and histones regulate this interaction. **A)** (Left) SEC of crosslinked CAF-1-PCNA complexes after DNA digestion using 1 μM PCNA, 0.15 μM RFC, 1.5 μM WT CAF-1 and 0.3 μM nicked pUC19. After crosslinking with 0.2% glutaraldehyde and quenching, the samples were treated with nuclease to digest the DNA plasmid. (Right) Coomassie SDS-PAGE of the collected fraction. The fractions that were used to prepare mass photometry samples are shown as Peak1 and Peak2. **B)** Mass photometry data of pooled fractions of Peak1 (left) and Peak2 (right) from experiment in panel A. Theoretical masses are listed and calculated masses from the fitted data are shown in each graph. Normalized counts are shown. Full length RFC is 220 kDa and the RFC-PCNA complex weigh 310 kDa; these RFC-containing particles are not observed in the data, as RFC is present at 10x lower concentration than CAF-1. **C)** SDS PAGE of protein-protein crosslinking reactions after DNA digestion. These reactions contain 50 nM fluorescently labeled PCNA, 15 nM full-length RFC, 15 nM pUC19 and increasing CAF-1 concentrations. PCNA fluorescence signal is shown. Full gels are shown in Supplementary Figure S3A. **D)** Quantification of the fluorescence intensity of bands in C. Data are shown as mean +/- SD of three independent experiments. **E)** SEC and SDS-PAGE of crosslinked CAF-1-PCNA complexes after DNA digestion with CAF-1_WHD*, as in panel A. WT curved is shown in dashed gray line for comparison. **F)** SDS PAGE of crosslinking reactions containing fluorescent PCNA (5.5 μM) and H3-H4 (H4-E63C, 1.5 μM dimer concentration), CAF-1 or tCAF-1 (1.5 μM). DNA or RFC is not present in these reactions. **G)** SDS PAGE of crosslinking reactions containing fluorescent PCNA (5.5 μM) and H3-H4 (H4-E63C, 1.5 μM dimer concentration), CAF-1 or CAF-1_ΔAD (1.5 μM). DNA or RFC is not present in these reactions.

To further evaluate the stoichiometry of CAF-1-PCNA-DNA complexes, we monitored complex formation using crosslinking at limiting concentrations of fluorescently labeled PCNA loaded onto DNA, which also allows us to estimate binding affinities. Without CAF-1, PCNA crosslinks with the clamp loader RFC (calculated molecular weight is 310 kDa) to form a complex that runs at the same height as 2xCAF-1-PCNA (Figure 3C and Supplementary Figure S3A). This band increases in intensity as we increase CAF-1 concentration above 100 nM while RFC is kept constant (Figure 3C-D and Supplementary Figure S3A). Interestingly, above 350 nM, we observed that 3xCAF-1-PCNA complexes formed while the 2xCAF-1-PCNA band became less pronounced (Figure 3C-D and Supplementary Figure S3A). We observed only a small fraction of 1xCAF-1-PCNA complexes (Figure 3CD and Supplementary Figure S3A) in line with the mass photometry results (Figure 3B). Together, these experiments demonstrate that CAF-1 prefers to bind PCNA on DNA with a 2:1 stoichiometry at concentrations around 100 nM. Above 350 nM, additional CAF-1 complexes can associate with DNA-loaded PCNA. Interestingly, only a very small fraction of CAF-1-PCNA complexes at a 1:1 stoichiometry is observed. This is in line with our previous hypothesis that two CAF-1 complexes cooperatively associate on DNA (54, 74), and it shows that it also applies to PCNA-dependent CAF-1 chromatin assembly.

To ask if this assembly is important for CAF-1 histone chaperone function, we tested if mutations in the WHD domain affected the stoichiometry of CAF-1-PCNA complexes. Indeed, the WHD is important for the cooperative DNA binding of CAF-1 and for its function in cells (54), and mutations in the WHD affect the PCNA-dependent nucleosome assembly activity of CAF-1 in the PCNA-NAQ assay (Figure 2C). CAF-1_WHD* affected the composition of CAF-1-PCNA complexes, with a reduction in the formation of 2:1 or 3:1 CAF-1-PCNA complexes in solution (Figure 3E). This explains why this complex is inactive in PCNA-dependent nucleosome assembly (Figure 2C) and argues that two CAF-1 complexes are required for histone deposition also in the context of PCNA.

### Histones further promote the CAF-1-PCNA interaction

Although histones are not strictly required for the formation of a CAF-1-PCNA complex on DNA (Figure 1), CAF-1 tightly binds H3-H4 during DNA replication. Thus, we set out to investigate if histones affect CAF-1 binding to PCNA. To this end, we investigated the role of histones on the CAF-1-PCNA interaction in the absence of DNA, because in DNA-containing reactions histones would be immediately deposited onto DNA, making it impossible to assess their effect on the CAF-1-PCNA interaction. As shown above, CAF-1 does not bind to PCNA when DNA is missing from the reaction (Supplementary Figure S1E). However, pre-incubation of CAF-1 with H3-H4 promotes the interaction between CAF-1 and PCNA in the absence of DNA in crosslinking experiments (Figure 3F). Deletion of the N-terminal region in Cac1, which contains the PIPs and KER domain (as in the truncated tCAF-1 construct, Supplementary Figure S1D), prevents the CAF-1-PCNA interaction (Figure 3F), confirming that this region is responsible for binding to PCNA within the complex. Interestingly, the interactions between CAF-1-H3-H4 and PCNA in the absence of DNA could not be observed from a SEC purification (Supplementary Figure S3B), suggesting that it is more dynamic than the interaction that is mediated by DNA.

Previous work has shown that H3-H4 binding to the CAF-1 acidic domain induces conformational changes at the PIPs, KER and WHD regions, that are important for CAF-1 histone chaperone function (54, 75). These conformational changes could be mimicked by deleting the acidic domain in CAF-1 (54), we thus generated a mutant carrying such deletion (CAF-1_ΔAD) to test if these conformational changes control the CAF-1-PCNA interaction. Strikingly, crosslinking between full-length CAF-1_ΔAD and PCNA shows efficient complex formation in absence of DNA and histones in crosslinking experiments (Figure 3G). Moreover, CAF-1_ΔAD efficiently forms complexes with PCNA on DNA at lower concentrations than WT CAF-1 (below 100 nM, Supplementary Figure S3C), suggesting an increase in binding affinity for this mutant to DNA-loaded PCNA. These data argue that changes that occur upon neutralization of the acidic domain (i.e. mimicking histone binding) in CAF-1 promote interactions with PCNA. Together these data suggest that histones are not required *per se* for PCNA binding on DNA, however they may promote the CAF-1-PCNA interaction via conformational changes that involve the N-terminal region in Cac1.

### CAF-1 inhibits DNA synthesis by Polε, but not Polδ, via PCNA

At replication forks PCNA binds several proteins, most prominently the replicative DNA polymerases on both daughter strands. As replicated DNA is readily assembled into chromatin at replication forks (22, 23), we next asked how DNA polymerases and CAF-1 may share or compete for binding to PCNA.

To this end, we first investigated the effects of CAF-1 on PCNA-mediated DNA synthesis by the leading- and lagging-strand DNA polymerases Polε and Polδ, in a primer extension assay. In this assay, the extension of a fluorescent DNA primer that is annealed to an RPA-coated single stranded plasmid is monitored over time. As previously shown, yeast Polδ and Polε efficiently synthesized DNA in a PCNA-dependent fashion with distinct kinetics (Supplementary Figure S4A) (15–17, 77). We found that adding CAF-1 had minimal effects on DNA synthesis by Polδ in this primer extension assay (Figure 4A). However, CAF-1 had a strong inhibitory effect on DNA synthesis by Polε, at concentrations of 150 nM where CAF-1 binds PCNA with a 2:1 stoichiometry (Figure 4B, 3D). This effect resolved at later time points and was dose-dependent, indicative of competitive inhibition (Figure 4B and Supplementary Figure S4B). This suggests a dynamic and steric effect of CAF-1 on Polε-mediated DNA synthesis. To test if this inhibition involved a crosstalk on PCNA, we used CAF-1 mutants that do not bind to DNA-loaded PCNA (i.e. CAF-1_PIP** and CAF-1_KER*). These mutants did not inhibit Polε activity (Figure 4CD), demonstrating that the inhibitory effect of CAF-1 on Polε is exerted via PCNA. These data are consistent with CAF-1 and Polε competing for binding on PCNA.

**Figure 4:**
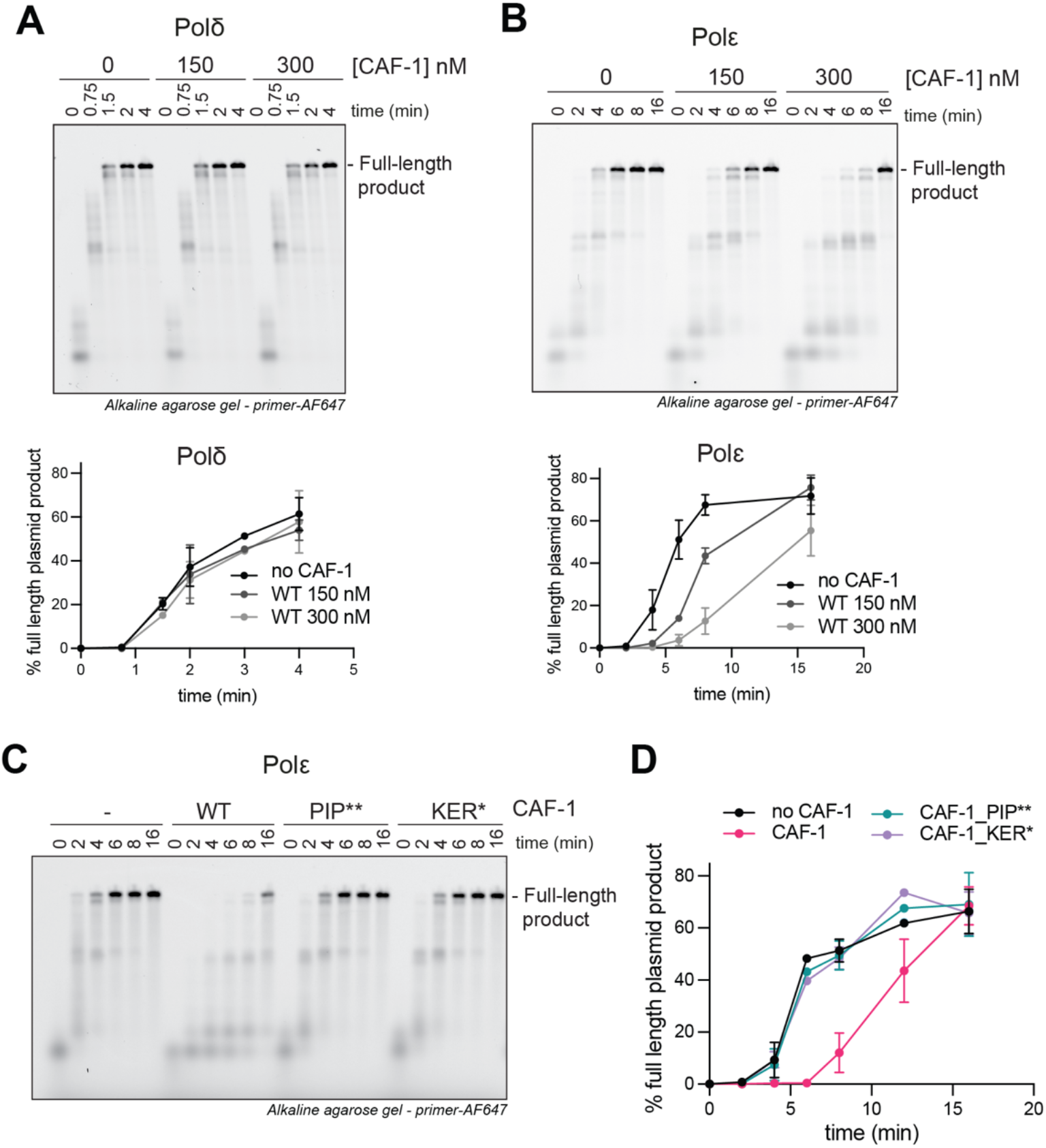
CAF-1 competes with Polε, not with Polδ, for PCNA binding. **A-B)** (Top) Fluorescence scan of a denaturing alkaline agarose gel of primer extension reactions with Polδ (A) or Polε (B). The fluorescently labelled primer signal is shown. The polymerases were at 120 nM, PCNA 480 nM and CAF-1 concentrations as shown. (Bottom) Quantification of the full-length product band relative to the total fluorescence in each lane (expressed as percentages) from the top panels. Mean ±SD are shown for independent experiments (Polδn=2 - Polε n=4). **C)** Fluorescence scan of denaturing alkaline agarose gel of primer extension reactions with Polε withCAF-1_PIP** and CAF-1_KER* mutants (300 nM). **D)** Quantification of primer extension by Polε as in Figure A and B.

In contrast to Polε, our data suggest that CAF-1 and Polδ efficiently share PCNA (Figure 4A). Previous studies have shown that Polδ has a tighter binding affinity for PCNA (Kd_app_=13,7 nM) than Polε (Kd_app_=326 nM) (17). We found that CAF-1 binds PCNA on DNA with intermediate binding affinity (^~^100 nM) (Figure 3C-D). Therefore, we tested whether Polδ might simply outcompete CAF-1 on PCNA, unlike Polε. To this end, we used the CAF-1_ΔAD mutant which shows tighter binding affinities for DNA-loaded PCNA (estimated <50 nM, in the same range as Polδ) (Supplementary Figure S3C). While inhibiting Polε even more strongly than WT CAF-1, this mutant still had only a minor effect on Polδ activity when added at 300 nM (Supplementary Figure S4C). Therefore, differences in PCNA binding affinities between the two polymerases do not solely explain the differences in their crosstalk to CAF-1, suggesting that steric effects might play a role. This is also supported by the limited interface that is used by Polδ in binding DNA-loaded PCNA in recent cryo-EM structures (Supplementary Figure S4D) (78, 79). Together, these results support a model in which CAF-1-mediated nucleosome assembly is differentially linked to DNA synthesis on the two daughter strands.

### Polε function and interplay with CAF-1 are independent of histone binding

During DNA replication in cells, Polε and CAF-1 both bind H3-H4 (72, 80, 81). Thus, we set out to test whether histones regulate the crosstalk between CAF-1 and the DNA polymerases on PCNA.

First, we used fluorescence polarization assays to determine the binding affinity of Polε for H3-H4 and find that Polε binds H3-H4 with a Kd=28 nM. This is a 25 times lower affinity than that of CAF-1 (Kd=1,1 nM) (Supplementary Figure S5A). Nevertheless, this would be sufficient to efficiently bind histones in our assay, where Polε is present at 120 nM. Polδ has background binding to H3-H4 (Kd≥300 nM), similarly to RPA which is also present in the reactions (Kd≥300 nM) (Supplementary Figure S5A). These data show that Polε and CAF-1 efficiently bind H3-H4, while Polδ and RPA do not bind histones in our assays.

To test the effect of histone binding in the crosstalk between CAF-1 and the DNA polymerases, we pre-incubated either the DNA polymerase or CAF-1 with H3-H4 and monitored how this affected DNA synthesis in primer extension assays. Polε activity was not affected by the addition of H3-H4 (Figure 5A) and the CAF-1-dependent inhibition of Polε was also largely unaffected by the presence of H3-H4 (Figure 5A). As expected, the addition of histones to reactions containing Polδ had no effect on DNA synthesis or on its crosstalk to CAF-1 (Figure 5B). These data demonstrate that histones do not alter the differential effects that CAF-1 has on Polδ and Polε via PCNA, arguing that this interplay is relevant during chromatin assembly at replication forks when histones are bound to the histone chaperones.

**Figure 5:**
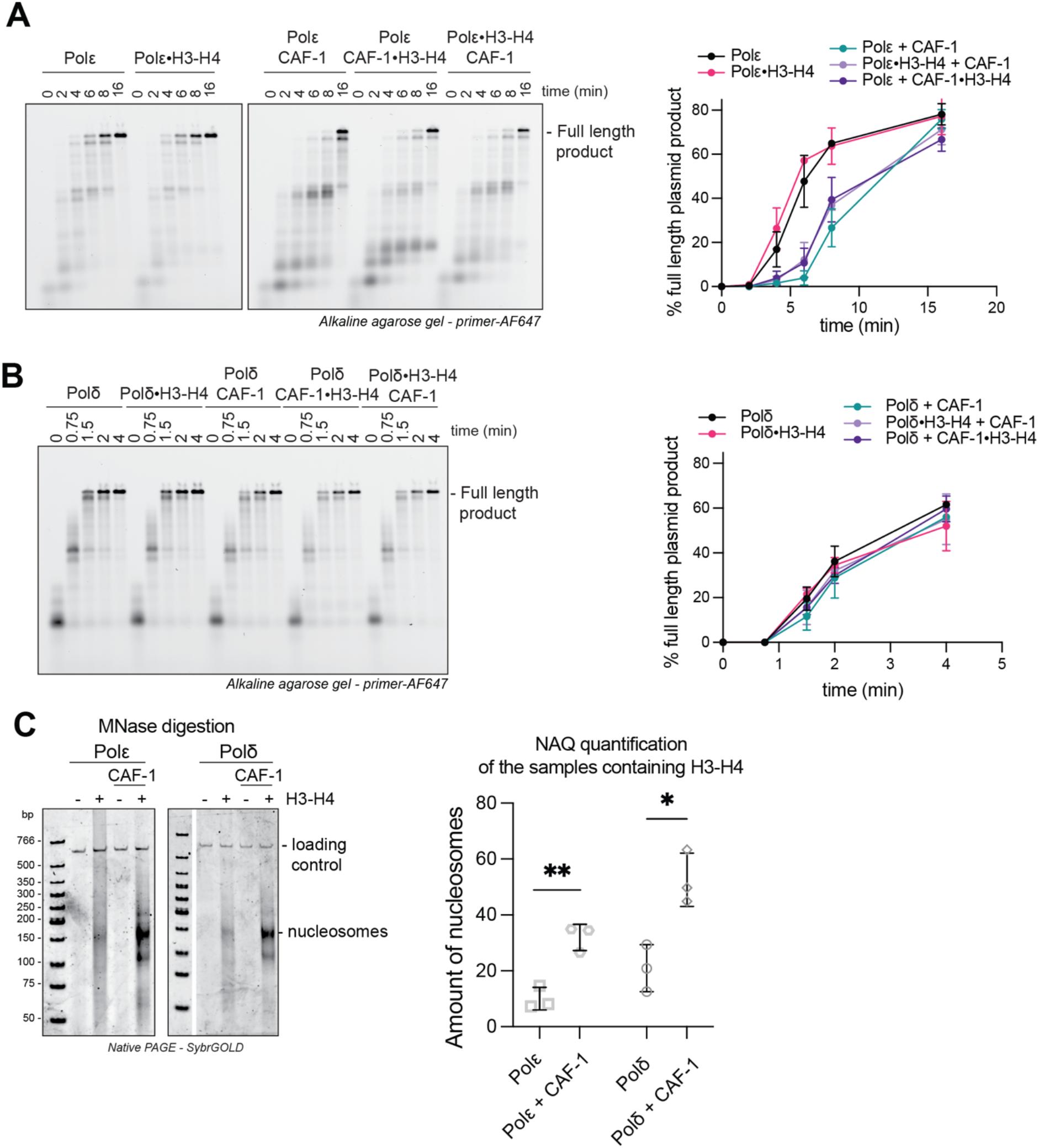
Polε function and interplay with CAF-1 are independent of histone binding. **A-B)** (Left) Fluorescence scan of denaturing alkaline agarose gel of primer extension reactions with Polε (A) or Polδ (B) in the presence of H3-H4. H3-H4 were either preincubated with the DNA polymerase or with CAF-1, as indicated by the •. The fluorescently labelled primer signal is shown. (Right). **C)** (Left) Native PAGE stained with SybrGOLD to detect protected DNA fragments following MNase digestion during primer extension reactions with Polε or Polδ in presence of CAF-1. H3-H4 were co-incubated with the polymerase or with CAF-1 throughout the reaction, H2A-H2B were added at 16min and samples were immediately treated with 80 units MNase. (Right) Bioanalyzer-based quantification of protected nucleosomal fragments from samples on the left, relative to the loading control band in each lane. Mean ± SD is shown for three independent experiments. * p<0.05, ** p>0.01 (unpaired t-test comparing Polε or Polδ to the condition containing CAF-1).

Previous studies have shown that CAF-1, Polε or RPA assemble chromatin during DNA replication (54, 80–82). As these proteins are all present in our assays, we set out to directly test which of these histone chaperone can assemble nucleosomes in these reconstitutions. To this end, we combined primer extension reactions with NAQ-based readouts to measure histone deposition (i.e. nucleosome assembly). Because in this assay the substrate is RPA-coated single-stranded DNA, nucleosome formation occurs only after DNA synthesis. In the absence of CAF-1, Polε containing reactions show background levels of nucleosome assembly (Figure 5C). These levels are even lower than the histone deposition that we observe with Polδ, which we used as a negative control because it can synthesize DNA but does not bind histones (Figure 5C). Both reactions contain RPA, indicating that this complex also does not stimulate nucleosome assembly. However, the addition of CAF-1 strongly increases nucleosome assembly in both conditions (Figure 5C). Similar results were observed when we measure nucleosome assembly on double-stranded DNA fragments using each histone chaperone complex in isolation (Supplementary Figure S5B), which shows that only CAF-1 can stimulate nucleosome assembly. Together, our data demonstrates that Polε and RPA are not intrinsically capable of nucleosome assembly in a replication-coupled manner, suggesting the main histone assembly factor in these reconstitutions is CAF-1.

### CAF-1 deposits newly synthesized H3-H4 on both daughter strands in cells

We identified a differential crosstalk of CAF-1 with Polε and Polδ,likely through their differential interaction with PCNA. As CAF-1 and Polε compete for binding on PCNA, we wondered whether CAF-1 is able to assemble nucleosomes on the leading strand. To address this question directly in cells, we used mouse embryonic stem cells (mESCs) and employed Sister Chromatid after Replication Sequencing (SCAR-seq) (67, 68). This is a genomic method that measures relative protein abundance on the two newly replicated daughter DNA strands, which allowed us to investigate whether depletion of CAF-1 results in a bias in deposition of new histones towards the leading strand.

We first generated a mESC line expressing a CAF-1 p150 subunit that is N-terminally tagged with FKBP12 (named dTAG-*Chaf1a*). dTAG-*Chaf1a* is targeted for proteasomal degradation in the presence of the degrader compound dTAG (83). In these cells, CAF-1 p150 is degraded within 1-2 hour of dTAG treatment (Supplementary Figure S6A), allowing acute depletion of CAF-1 during DNA replication to study its function with minimal pleiotropic effects. We observed that CAF-1 degradation led to a marked reduction of new histones, identified by H4 unmethylated at lysine 20 (H4K20me0), and DNA synthesis, recapitulating known effects of CAF-1 insufficiency (Supplementary Figure S6B-E)(36, 37, 84, 85).

Parental H3-H4 are recycled in a quasi-symmetrical fashion at replication forks, where each newly replicated DNA strand receives about 50% of these histones (67, 81). Simultaneously, newly synthesized histones are also symmetrically assembled on the two daughter strands to maintain nucleosome density on replicated DNA (67, 81). Control SCAR-seq experiments in untreated dTAG-*Chaf1a* mESCs confirmed these observations, using H3K27me3 as a marker of parental histones (24) and H4K20me0 to mark new histones (86, 87) (Figure 6A). Upon dTAG treatment, the total reads in the EdU inputs decreased, consistent with reduced DNA synthesis (Supplementary Figure S6F). Moreover, we observed a 2-fold reduction in reads for the H4K20me0 pulldown upon CAF-1 depletion (Figure 6B), with the H3K27me3-marked parental histones showing a comparable increase (Figure 6B). This could be due to increased MNase accessibility or to effects on parental histones dynamics. This demonstrates that CAF-1 is required for deposition of newly synthesized histones, while parental histone recycling occurs independently of CAF-1. Consistently, parental histones were distributed nearly symmetrically to both daughter strands in the absence of CAF-1 (Figure 6A). Moreover, depletion of CAF-1 did not result in an asymmetric distribution of the new histones that were deposited in this context. This argues that CAF-1 is active on both the leading and lagging strands of active replication forks in mESCs, as are backup systems such as HIRA-dependent gap filling (88).

**Figure 6:**
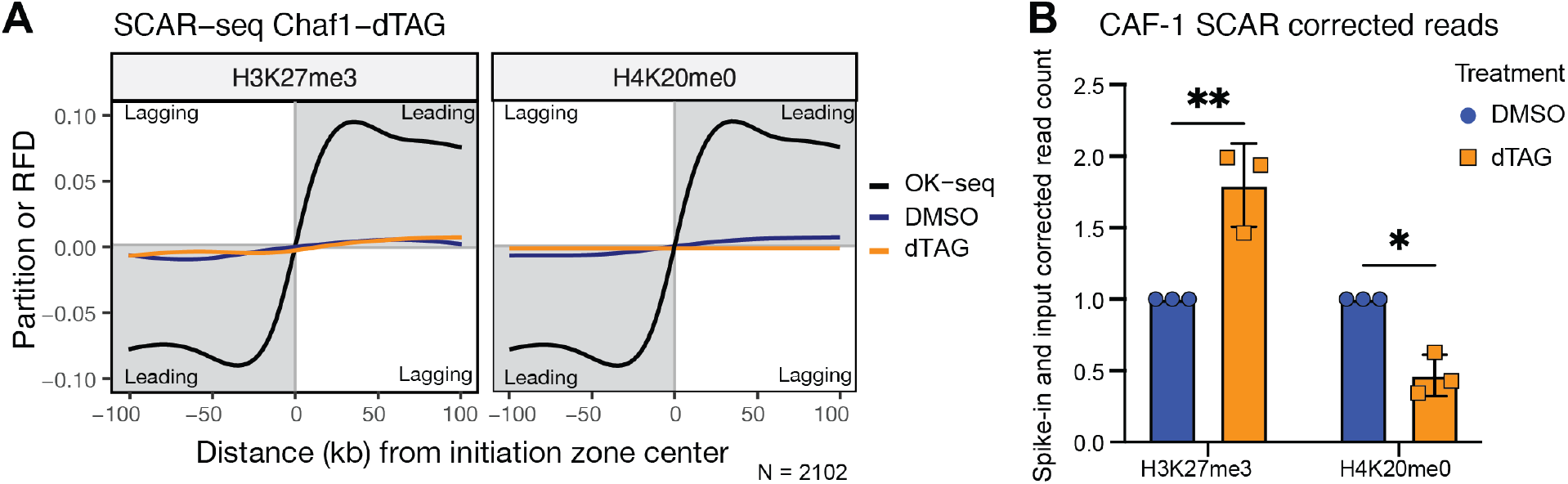
CAF-1 deposits newly synthesized H3-H4 on both leading and lagging strands. **A)** Average SCAR-Seq profile of parental (H3K27me3) (left) or newly synthesized (H4K20me0) (right) histones across all replication initiation zones (N(IZ) = 2102) in control (DMSO) or dTAG treated samples. Partition is calculated as the proportion of forward (F) and reverse (R) read counts. Replication fork directionality (RFD) in WT cells measured by Okazaki fragment sequencing (OK-Seq) is shown for comparison. **B)** Spike-in normalized values for parental (H3K27me3) and new (H4K20me0) histone modification shows a significant reduction in H4K20me0 samples when CAF-1 is depleted. n=3 independent experiments. * p<0.05, ** p<0.01 (two-way ANOVA).

Together, these data show that CAF-1 functions on both the leading and lagging strand of replication forks in mESCs, where it primarily deposits newly synthesized histones. CAF-1 removal affects the incorporation of these histones on both daughter strands equally without challenging parental histone recycling. This indicates that although CAF-1 and Polε compete for PCNA, both machineries efficiently function on the leading strand.

### CAF-1 and Polε compete for PCNA within the replisome

As both CAF-1 and Polε function on the leading strand, we used biochemical reconstitutions to investigate the role of replisome proteins in the interplay between CAF-1 and Polε. Polε is an integral and essential component of the CMG complex at replication forks (10, 17, 89–91). We purified the yeast replisome components that were previously shown to recapitulate physiological DNA replication *in vitro* (8, 9) (Supplementary Figure S7A). Our preparations are active as they promote replication of ARS1-containing DNA plasmids in a manner that depends on the presence of the Dbf4-dependent kinase (DDK) (Supplementary Figure S7B) (9).

To focus on Polε activity, we used a pulse-chase setup in which we omitted Polδ. This allowed us to quantify replication rates of the leading strand only (9). In this assay, Polε is capable of DNA synthesis in the absence of PCNA with a rate of ^~^0.47 kb/min (Figure 7A-B) (9). The addition of PCNA and its loader RFC to these reactions increases the rates to ^~^1 kb/min, recapitulating physiological speeds (Figure 7A-B) (9). Strikingly, the addition of CAF-1 led to a reduction in the rate to ^~^0,75 kb/min (Figure 7A-B), suggesting an inhibitory effect of CAF-1 towards Polε in the context of an active replisome. Consistent with this conclusion, the CAF-1_PIP** mutant did not reduce the speed of DNA replication (Figure 7A-B). This argues that a competition between CAF-1 and Polε on PCNA might occur at physiological replication forks.

**Figure 7:**
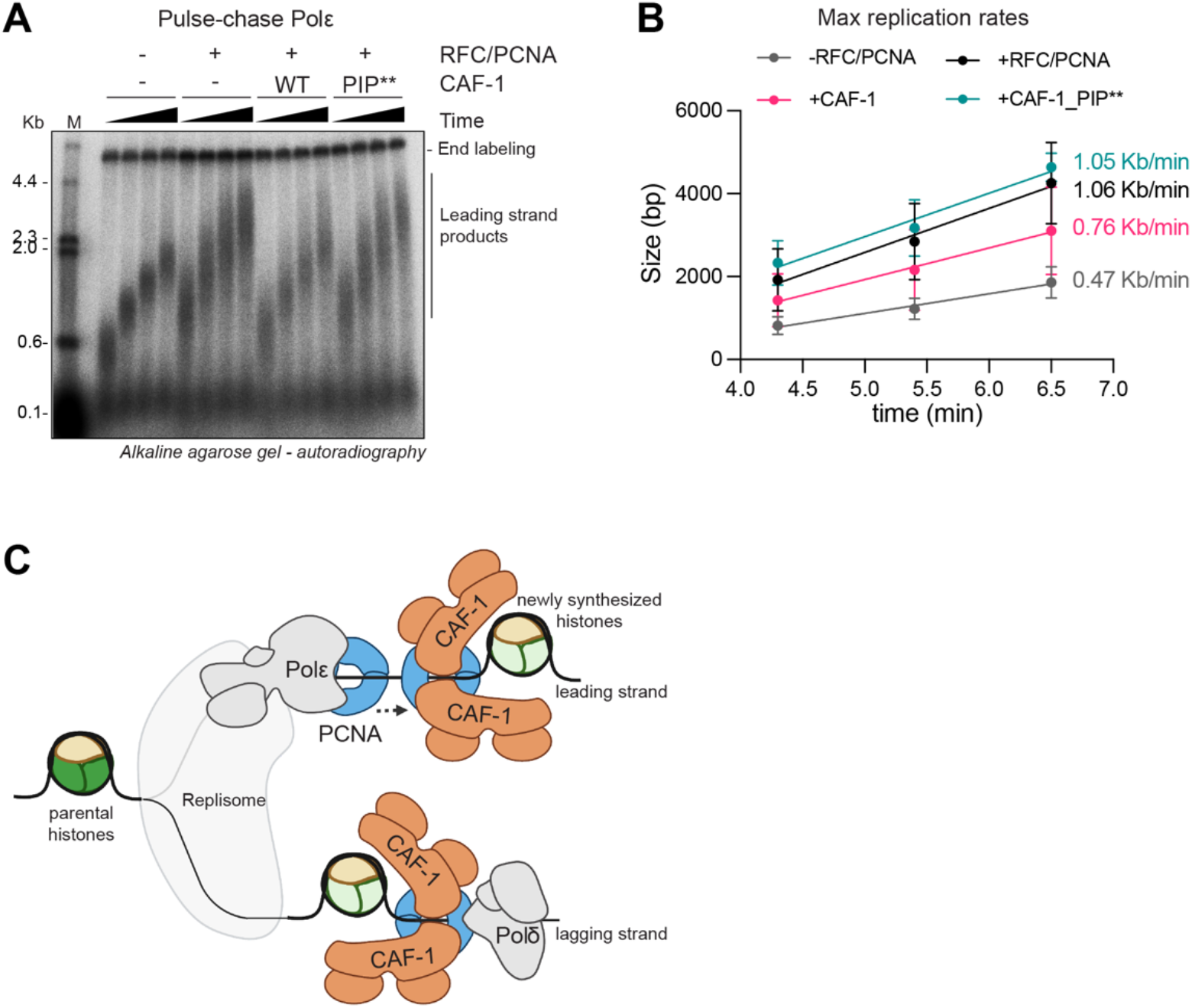
CAF-1 and Polε compete for PCNA binding within the replisome. **A)** Autoradiography scan of a denaturing agarose gel of DNA replication products from a pulse-chase experiment in presence of the yeast replisome (Polδ and TopoII are omitted). All proteins were present during the pulse step (3 minutes 20 seconds). After addition to the chase solution, reactions were stopped at the indicated time points (4.3, 5.4, 6.5 and 7.5 minutes). **B)** Quantification of the maximum replication fork speed for pulse-chase experiments in A. The length of the leading strand products was determined using ImageQuant. Product sizes were then plotted against time. The maximum fork speed was obtained by fitting the data to a linear regression. Data are shown as mean +/- SD of 3 independent experiments, with the exception of the CAF-1_PIP** where two independent experiments are included. **C)** The crosstalk of CAF-1 mediated nucleosome assembly with the DNA replication machinery differs between the leading and lagging strand of replication forks. Two CAF-1 complexes associate with PCNA on DNA to assemble a nucleosome. CAF-1 competes with the leading strand DNA polymerase Polε for PCNA binding, but not with the lagging strand polymerase Polδ. Nevertheless, CAF-1 deposits newly synthesized histones on both daughter strands. This means that on the leading strand, chromatin assembly by CAF-1 cannot occur simultaneously with DNA synthesis, while on the lagging strand both processes are coupled. The model was created with BioRender.com.

These data, combined with the observation that CAF-1 acts on both daughter strands (Figure 6A), support a model in which CAF-1-mediated nucleosome assembly on the leading strand is spatially separated from DNA synthesis. We propose that either, CAF-1 and Polε interact with distinct PCNA clamps, or that these two machineries alternate in binding to PCNA in a dynamic manner. In contrast, on the lagging strand there is a closer link between CAF-1 and Polδ, which can efficiently share PCNA. These observations suggest that distinct mechanisms act on the two daughter strands to couple DNA synthesis with CAF-1 mediated nucleosome assembly (Figure 7C).

## DISCUSSION

Our work provides insights into how the essential histone chaperone CAF-1 functions during genome replication. We show that CAF-1 recruitment and PCNA-dependent nucleosome assembly activity are regulated by a complex set of interactions between CAF-1 and PCNA, DNA and histones. Our results argue that several structural transitions regulate CAF-1, and we anticipate that these control the timing of CAF-1 arrival to, action on and departure from replication forks. This is linked with the interplay between CAF-1, PCNA and the DNA polymerases. CAF-1 competes with Polε for PCNA binding, while it can efficiently share PCNA with Polδ. Yet, CAF-1 deposits newly synthesized histones equally on both strands in mESCs. This competition between CAF-1 and Polε appears to be an integrated part of coordinating replication and nucleosome assembly and does not limit CAF-1 function. Our work thus implies that different mechanisms are in place on the leading and lagging strand to couple DNA and chromatin replication.

### Coupling of DNA synthesis and chromatin assembly on the two daughter strands

We propose that on the leading strand, CAF-1 and Polε either alternate in interacting with PCNA or bind distinct PCNA clamps. An alternation between Polε and CAF-1 is consistent with their binding properties to PCNA (15, 92), the competitive inhibition that we observe (Figure 4B), and the dependence of CAF-1 recruitment on sufficient DNA length at PCNA (Figure 1G). This interplay may further represent a mechanism for enabling continued PCNA loading on the leading strand during elongation (19) and it could directly affect replication speed *in vivo*. Both models evoke the need for regulatory steps in the loading of PCNA on the leading strand, where the CTF18 clamp loader may well play a role (19, 93). Either way, our data show that DNA synthesis and nucleosome assembly cannot occur simultaneously on the same PCNA ring, proving that these two processes may be functionally linked but they are physically separated on the leading strand. As Polε is a histone chaperone for parental H3-H4 during DNA replication in cells (80, 81), and as we show that CAF-1 primarily deposits newly synthesized histones (Figure 6B), the competition between CAF-1 and Polε on PCNA may represent a mechanism to control parental and new histone incorporation on replicated DNA. Interestingly, we did not detect histone deposition activity by Polε in our assays, indicating that additional factors may be functioning on the leading strand to promote parental nucleosome assembly. Our studies pave the way for future investigations in understanding how parental and new histone deposition pathways are integrated during DNA replication.

On the lagging strand, CAF-1 and Polδ may interact with PCNA simultaneously. This is consistent with structural data showing that Polδ occupies only one of the PCNA monomers on the DNA-loaded clamp (78, 79) (Supplementary Figure S4D). Moreover, this is in line with cellular evidence that closely links Okazaki fragment size with nucleosome assembly by CAF-1 (54, 94). Finally, this aligns with Polδ not having histone chaperone activity (Supplementary Figure S5A), thus requiring CAF-1 to be in close proximity to readily assemble chromatin on newly synthesized DNA. In this model, it remains unclear how histone chaperones that bind parental H3-H4 intercalate with CAF-1 to deposit recycled histones. MCM2 plays an important role in parental histone dynamics on the lagging strand (67), but it is unlikely to be responsible for histone deposition due to its localization ahead of the fork. RPA is a good candidate for nucleosome assembly on the lagging strand (82) but we observe no activity for this complex in our primer extension assays. In our view, DNA polymerase alpha (Polα) is an attractive player (95–97), but it remains unclear how this can work in relation to PCNA loading and Polδ function. Our reconstitutions provide suitable systems to test these hypotheses in the future.

In summary, our work investigates how the replisome and histone chaperones integrate genome and chromatin replication and reveals mechanistic differences in how CAF-1 shares PCNA with the two replicative DNA polymerases. We propose that the observed differences in CAF-1 mediated deposition of newly synthesized histones have evolved to accommodate the distinct parental histone recycling mechanisms on the two daughter strands and the inherent asymmetry of DNA replication.

## MATERIAL AND METHODS

### Protein expression and purification

CAF-1 and PCNA mutants were made using standard mutagenesis procedures and purified following the wildtype purification protocols. We used yeast proteins, with the exception of *X.laevis* histones. Several proteins used in our study were expressed and purified as previously described. This includes PCNA (56), Polδ and Polε (8, 9), CAF-1 and its mutants (54). Lyophilized *Xenopus laevis* histones were purchased from the Histone Source at CSU, Fort Collins, CO, USA. These were labeled with maleimide dyes (when required) and refolded as in (57, 58). ORC, cdc6, Mcm2/7-cdt1, DDK, cdc45, Dpb11, GINS, S-CDK, Mcm10, RPA, Polα, Ctf4, Sld3/7, Sld2 and TopoII were purified as in (8). Csm3/Tof1, Topo I, RFC and PCNA were purified as described in (9). Mrc1 was expressed and purified following the procedure described in (59). PCNAK164C contains all native cysteines with the addition of an engineered K164C mutation that adds an extra cysteine on the exposed protein surface. All the concentrations for PCNA reported here refer to the monomer concentration. Additional purification protocols are:

#### RFCΔN (used in Figures 1–3A,B,E-G,4)

Rosetta2 cells containing pBL481-RFCΔN (60) were grown in 4 liters of Terrific Broth at 37°C to A_600_=1.6. The temperature was shifted to 18°C and cells were incubated for 30 more minutes before adding 0.3 mM isopropyl-thiogalactoside (IPTG). The expression was incubated for 16 hours and cells were harvested and resuspended in 30 mM HEPES pH 7.5, 0,5 mM EDTA, 10% Glycerol, 200 mM NaCl, 1mM DTT, 0,5 mM p-methylphenyl-sulfonyl fluoride (PMSF) in presence of COMPLETE EDTA-free protease inhibitor (Roche). Cells were lysed with sonication. DNA was precipitated with 0.5% of poly(ethyleneimine) (PEI) and the lysate was clarified by centrifugation for 30 minutes at 21000xg. RFCΔN was precipitated with 0.28 g/ml of AmSO4. The precipitates were collected by centrifugation at 12000xg for 45 minutes. Pellets were resuspended in 50ml of 30 mM HEPES pH 7.5, 0,5 mM EDTA, 10% Glycerol + 1mM DTT, 0.5mM PMSF in presence of COMPLETE EDTA-free protease inhibitor. The lysate was next dialyzed (12-14 MWCO) against 2 liters of 30 mM HEPES pH 7.5, 0,5 mM EDTA, 10% glycerol + 100 mM NaCl, 1mM DTT for 2 hours. RFCΔN was injected on HiTrap SP HP 5ml column (Cytiva) equilibrated in buffer SP-A (30 mM HEPES pH 7.5, 10% Glycerol, 100 mM NaCl, 1mM TCEP). The column was washed with 25ml of SP-A buffer and eluted in a gradient of SP-B buffer (30 mM HEPES pH 7.5, 10% Glycerol, 800 mM NaCl, 1mM TCEP) along 60ml. Fractions containing RFCΔN were analyzed by SDS-PAGE and were pooled together. RFCΔN was then mixed with 3 ml of nickel beads equilibrated in His-A buffer (20mM HEPES 7.5, 200mM NaCl, 20 mM Imidazole, 10% Glycerol, 1 mM TCEP). Beads were washed with 100 ml of His-A buffer and RFCΔN was eluted with His-B buffer (20mM HEPES 7.5, 300mM NaCl, 300 mM Imidazole, 10% Glycerol, 0.05% ampholytes, 1 mM TCEP). Fractions containing RFC were concentrated and further purified on HiLoad 16/600 Superdex 200 in 20mM HEPES 7.5, 200mM NaCl, 10% Glycerol, 1 mM TCEP, 0.05% ampholytes. RFCΔN was concentrated and stored at −80°C.

#### RPA from bacterial expression (used in Figure 4–5)

Rosetta2 cells transformed with pRSF-Duet, RPA, a gift from Xiaodong Zhang (61), were grown in 2 liters of Terrific Broth at 37°C for 16 hours until A600=1.8. Cells were placed at 25°C and RPA expression was induced with 0.3 mM IPTG for 3 hours. Cells were harvested and resuspended in lysis buffer (50 mM HEPES pH 7.5, 500 mM NaCl, 5% glycerol, 0.01% IGEPAL CA-630, 1mM TCEP) in presence of COMPLETE EDTA-free protease inhibitor (Roche). Cells were sonicated and the lysate was clarified by centrifugation at 50000xg for 50 minutes. The supernatant was recovered and injected on HisTrap HP 5ml column equilibrated in lysis buffer. The column was washed with 50 ml of lysis buffer, 100 ml of His-A buffer (50 mM HEPES pH 7.5, 750 mM NaCl, 5% glycerol, 0.01% IGEPAL CA-630, 1mM TCEP, 30 mM Imidazole), and 25 ml of lysis buffer respectively. RPA was then eluted in a gradient of His-B buffer (50 mM HEPES pH 7.5, 150 mM NaCl, 5% glycerol, 0.01% IGEPAL CA-630, 1mM TCEP, 250 mM Imidazole) along 50ml. Fractions containing RPA were pooled and diluted in 50 mM HEPES pH 7.5, 5% glycerol, 0.01% IGEPAL CA-630, 1mM TCEP to bring the salt concentration to 150 mM NaCl. RPA was next injected on HiTrap Heparin HP 1ml equilibrated in QA buffer (50 mM HEPES pH 7.5, 150 mM NaCl, 5% glycerol, 0.01% IGEPAL CA-630, 1mM TCEP). The column was washed with 20 ml of QA buffer and RPA was eluted in a gradient of QB buffer (50 mM HEPES pH 7.5, 1000 mM NaCl, 5% glycerol, 0.01% IGEPAL CA-630, 1mM TCEP) along 40ml. Fractions containing RPA were pooled together and injected on HiLoad 16/600 Superdex 200 and eluted in 50 mM HEPES pH 7.5, 300 mM NaCl, 5% glycerol, 0.01% IGEPAL CA-630, 1mM TCEP. RPA was concentrated and stored at −80°C.

### Protein labelling with fluorescent dyes

Histones H2A-H2B (containing H2B-T112C) and H3-H4 (containing H4-E63C) were labeled with maleimide AlexaFluor-647 (AF647) or AlexaFluor-488 (AF488) respectively (57, 58), as indicated.

PCNAK164C (containing all native cysteines in addition to the engineered K164C) was used with Alexa Fluor 546. PCNAK164C was diluted in labelling buffer (50 mM MOPS pH 7.0, 125 mM NaCl, 5 mM NaAc, 1 mM EDTA, 1 mM TCEP) to a final concentration of 1mg/ml. A 10-fold excess of TCEP was added to PCNA to ensure that all cysteines are effectively reduced. PCNA was then incubated with a 10-fold excess of AlexaFluor546. The reaction was incubated for 2 hours at room temperature, then quenched with 20 mM DTT final concentration for 30 minutes. Labelled PCNA was then concentrated and injected on a Superdex 75 increase 10/300 column to remove free dye. PCNA was eluted in 20 mM HEPES, 125 mM NaCl, 1 mM TCEP. Fractions containing labelled PCNA were pooled and concentrated, and the protein was stored at −80°C.

### Annealing of linear DNA fragments

Single-stranded DNA oligos of different lengths were purchased from IDT, either desalted (unlabeled oligos) or HPLC-purified (Alexa Fluor 647-conjugated oligos). For each length (18mer, 33mer, 43mer, 53mer) a forward oligo and a reverse oligo in reverse complement sequence were ordered. The 18mer and 33mer forward oligos included a 5’ Alexa Fluor 647 label. Forward and corresponding reverse oligos were mixed in a 1:1 stoichiometric ratio at 20 μM each (18mer and 33mer) or 40 μM each (43mer and 53mer) with a final of 20 mM HEPES pH 7.5 and 25 mM NaCl. The mixed oligos were annealed by heating up to 95 C for 3 min, and then slowly cooled to room temperature over several hours. Annealed DNA was stored at −20 C.

### EMSA

Native DNA-protein complexes were allowed to form in NA buffer: 25 mM TRIS pH 7.5, 150 mM NaCl, 1 mM EDTA, 0.02% Tween-20, 1 mM TCEP. Increasing amounts of CAF-1 (0 to 5 μM) were incubated in buffer for 10 min before addition of DNA (50 nM). Single stranded DNA oligos for 18bp, 33bp, 43 or 53 bp were purchased from IDT (labelled with AlexFluor647 at their 5’ end, Supplementary Table S1) and annealed prior to the EMSA experiments. 10% final concentration of glycerol was added before loading the samples into a 6% PAGE. Gels were scanned for fluorescence and then stained with SybrGOLD before imaging with Amersham Image Quant 800. The data was analyzed and plotted using FIJI and GraphPad Prism. We quantified the fluorescent signal of the unbound DNA band. We calculate the percentage of unbound DNA relative to the no CAF-1 condition. %bound DNA is then expressed as 100 - percentage of unbound DNA. The Kd-values were calculated using a one site binding curve with hill slope in GraphPad Prism. The 18bp data was fitted to a one site binding curve with a Hill coefficient constrained to 1.

### PCNA-CAF-1 binding experiments on SEC

We used pUC19 plasmid as DNA template for PCNA loading. This plasmid was nicked using the restriction enzyme Nt.BspQI for 8 hours at 50°C, and was subsequently purified via phenol chloroform extraction. Reactions were performed in PCNA loading buffer 50mM HEPES pH 7.5, 200 mM NaCl, 0.01% IGEPAL CA-360, 1mM TCEP. PCNA (30 μM) was incubated for 5 minutes at 30°C with nicked pUC19 (0.3 μM) and RFCΔN (0.5 μM), in the presence of MgCl_2_ (10 mM) and ATP (3 mM). Next, CAF-1 (5μM) was added to these reactions and incubated for 15 minutes at room temperature. Samples were next spun down for 5min at 17000xg before injection on Superose 6 increase 3.2-300 columns connected to an AKTA pure system fitted with PEEK I.D. 0.25 mm tubing. Fractions were analyzed on 4-12% gradient SDS-PAGE run in MES buffer.

### PCNA-NAQ assay

We used pRC1765 (Addgene #141346, a gift from Rafael Fernández Leiro) as template for PCNA loading and nucleosome assembly. pRC1765 was nicked using the restriction enzyme Nt.BbvCI for 6 hours at 37°C, and was subsequently purified via phenol chloroform extraction. PCNA was loaded on DNA in PCNA loading buffer, in a final volume of 11μl: PCNA (10.9 μM) was added to an equimolar mixture of nicked and supercoiled pRC1765 (47.3 nM each), RFCΔN (1.1μM) in presence of MgCl_2_ (8 mM) and ATP (10.9 mM). This reaction was incubated at 30°C for 5 minutes. First, samples were diluted with 25μl of NA buffer in order to decrease the high concentration of MgCl_2_ which hinder proper nucleosome assembly, followed by addition of CAF-1•H3-H4 (0.1 μM final concentration for each – H3-H4 dimer concentration) to a final volume of 40 μl total. This tetrasome assembly step is incubated at room temperature for 15 minutes. Then, we added fluorescently labelled H2A-H2B dimers and incubated for 15 minutes at room temperature, to complete nucleosome formation (62). Samples were spin down for 5 minutes at 17000xg. 1μl of each reaction was mixed with 5μl of NA buffer and 5% sucrose final concentration for loading on 0.8% agarose gel and run for 90 minutes in 1X TAE (TRIS-Acetate EDTA) at 90 volts. 25μl of each reaction was digested with 80 units of MNase in a total volume of 100 μl (containing 50 mM TRIS pH 7.9, 5 mM CaCl_2_) at 37°C for 10 minutes. MNase was inactivated by addition of EDTA. A 621bp DNA fragment was added as loading control and the DNA was further purified as in (62). MNase-digested samples were loaded on 6% PAGE and stained with SybrGOLD. The data was analyzed and plotted using FIJI, and GraphPad Prism. The PCNA-mediated activity of CAF-1 is quantified as the percentage of fluorescence on nicked plasmid relative to the total intensity (nicked + supercoiled) for each condition.

### MNase-seq of PCNA-NAQ assay

MNase-seq was used to quantify nucleosome assembly in the PCNA-NAQ assay. In order to distinguish nicked and supercoiled DNA, we used two plasmids with different sequences: pRS415 and pLox3 (Supplementary Table S2). After MNase inactivation a 207 bp DNA fragment was added as loading control in these experiments. Purified MNase-digested products (containing the loading control DNA) were used to prepare a Illumina sequencing library. First, samples were purified using the CleanNGS kit (GC biotech #CNGS-0008), according to the manufacturer’s protocol. Next, the CleanNGS elute was adjusted to 25ul with 10mM Tris pH 7.5 and the ends of the digested DNA were repaired and phosphorylated at their 5’ end using the End-It DNA End-repair kit (Lucigen #ER0720). DNA was purified using MinElute PCR Purification Kit (QIAGEN #28006). Next, 3’A overhang were added to each fragment using the Klenow fragment (NEB #M0212M) and DNA was purified using MinElute PCR Purification Kit (QIAGEN #28006). Next, unique indexed DNA adapters (Supplementary Table S3) were ligated overnight at room temperature T4 DNA ligases (NEB # M0202L) to all fragments with A-overhangs and DNA was purified using MinElute PCR Purification Kit (QIAGEN #28006). Finally, all samples were amplified by a 8-cycle PCR-program using Phusion High-Fidelity DNA Polymerase (NEB #M0530L) using primers 5’-TCGTCGGCAGCGTCAGATGTGTATAAGAGACAGCTCGGCATTCCTGCTGAACCGCTCTTCCGATCT-3’and 5’-GTCTCGTGGGCTCGGAGATGTGTATAAGAGACAGTACACTCTTTCCCTACACGACGCTCTTCCGATCT-3’, prior to a final clean up using the MinElute Purification Kit (QIAGEN #28006). Samples were pooled with a total concentration of 100ng. The library was submitted for paired-end Illumina 150bp PE sequencing at Macrogen (Amsterdam). fastaq files are uploaded to OSF (https://osf.io/2vd4z/?view_only=5ffa1e0b749445da9b22a11577f3d47f). PCNA-NAQ-seq analysis was performed using custom scripts (https://github.com/deLaatLab/PCNA-NAQ-seq). The sequence data was demultiplexed by extracting reads that contained the ligated adapter index in both read ends and trimmed by removal of the 5’ adapter sequence from the reads. Demultiplexed reads were mapped against the pLox3, pRS415 and loading control DNA sequences using BWA mem v0.7.17 and filtered using samtools with SAM flag 780 and mapping quality 60 and saved as bam files. The bam files were imported in R and fragments mapping to pLox3 and pRS415 with fragment lengths between 125 and 160bp were selected for further analysis. The percentage of reads mapping to the nicked plasmid was calculated based on the total amount of reads found on both nicked and supercoiled plasmids. For coverage analysis pLox3 and pRS415 fragments were normalized for the total number of fragments mapping to the loading control sequence.

### Primer extension assays

Experiments with Polε were performed in 25 mM HEPES-KOH pH 7.5, 150 mM potassium acetate, 8 mM magnesium acetate 1 mM TCEP, 1 mM ATP and 0.2 mg/ml BSA. Experiments with Polδ were performed in 25 mM TRIS-HCl pH 7.5, 150 mM NaCl, 8 mM MgCl_2_, 1 mM TCEP, 1 mM ATP and 0.2 mg/ml BSA. We used single stranded plasmid DNA as template for DNA synthesis, and it was produced as previously described(63).The concentrations reported here are for the final reaction that contains all components.

Single-strand pBluescript SK(-) (Supplementary Table S4) was incubated for 5min at 80°C with a 5x excess of a 15bp oligonucleotide and allowed to slowly cool down. The primer sequence is: G*G*G* T*T*C*GTGCACACA conjugated to an Alexa Fluor 647 dye at the 5’ end (* indicates nucleotides containing phosphorothioate bonds). The annealing reaction was coated with RPA (0.6 μM) for 5 minutes at 30°C. Next, PCNA (0.48 μM) was loaded in presence of RFCΔN (0.12 μM) for 5 minutes at 30°C on DNA (12nM). Polε or Polδ (0.12 μM) were primed onto the primer-template DNA in presence of dCTP, dGTP and dATP (75 μM of each) for 5 minutes at 30°C. Finally, dTTP (75 μM) was added to start the reaction. CAF-1 was also added at this step, at 300 nM unless stated otherwise in the figures. Reactions were quenched at various timepoints with 10 mM EDTA final concentration. Samples were mixed with 2% sucrose, 100 mM NaOH final concentrations and were loaded on denaturing alkaline 1.2% agarose gel. Gels were run for 16 hours at 40V, and imaged on a Typhoon. The data was analyzed and plotted using FIJI, and GraphPad Prism. DNA synthesis is quantified as the intensity of the full-length plasmid band relative to the total intensity in the entire lane.

For MNase analysis, 30 μL of primer extension reactions at the final time point (16 minutes for Polε and minutes for Polδ) were mixed with 80 U of MNase in a total of 100 μL (containing 50 mM TRIS pH 7.9, 5 mM CaCl_2_) at 37°C for 10 minutes. MNase was inactivated by addition of EDTA. A 621bp DNA fragment was added as loading control and the DNA was further purified as in (62). MNase-digested samples were loaded on 6% PAGE and stained with SybrGOLD and run on a Bioanalyzer (Agilent) using DNA High sensitivity chips. The bioanalyzer data was analyzed by normalizing the nucleosome band (140-150 bp) to the loading control at 621 bp within each lane, as in (62). Data was then plotted in excel and GraphPad Prism.

### In-solution crosslinking experiments

#### CAF-1-PCNA on nicked plasmid

To buffer containing 20 mM Hepes pH 7.5, 200 mM NaCl, 1 mM TCEP, the following components were added in order at room temperature: 10 mM MgCl2 (from 100 mM stock), 3 μM PCNA, 0.15 μM RFCΔN, 0.3 μM nicked (with Nt.BspQ1) pUC19 plasmid (from 1 μM stock), 1 mM ATP. This mixture was incubated at 30 °C for 5 min to increase the efficiency of PCNA loading onto DNA. Then, 1.5 μM CAF-1 was added and incubated for 10 min at room temperature. The total NaCl concentration during the loading reaction and after adding CAF-1, taking into account the contributions from each component, ranged between 100 and 110 mM. Samples were diluted 2-fold in buffer containing 20 mM Hepes pH 7.5, 100 mM NaCl, 1 mM TCEP, 0.02% IGEPAL CA-630, and incubated at room temperature for 10 min. Samples were subjected to chemical crosslinking by addition of a final concentration of 0.2% glutaraldehyde (from a 2.5% stock in water). The samples were incubated at room temperature for 20 min before quenching the crosslinker by addition of 100 mM Tris pH 7.5 (from 1 M stock). To release the crosslinked complexes from the DNA, 10% of the sample volume Pierce universal nuclease, diluted 1:20 in buffer containing 20 mM Hepes pH 7.5, 100 mM NaCl, 5 mM MgCl2, and 1 mM TCEP, was added. After incubation at room temperature for 10 min, 50 mM EDTA was added to quench the nuclease. Samples were spun down for 15 min at 13,000 xg at 4 °C and the supernatant was transferred to a new tube.

#### Complex formation of CAF-1 and PCNA on linear DNA

Linear DNA fragments with lengths of 18, 33, 43 or 53 bp were mixed with PCNA and CAF-1 in buffer containing 20 mM Hepes-KOH pH 7.6, 100 mM KCl, 0.01% IGEPAL CA-630 and 1 mM TCEP, and incubated on ice for 10 min. The final mixture contained 1.5 μM DNA, 4.5 μM Alexa Fluor 456 -labeled PCNA (concentration for a monomer), and 3 μM CAF-1. Samples were subjected to chemical crosslinking by diluting 3-fold in the same buffer and addition of a final concentration of 0.2% glutaraldehyde (from a 2.5% stock in water). The samples were incubated at room temperature for 20 min before quenching the crosslinker by addition of 100 mM Tris (from a 25x TAE stock containing 1 M Tris). Samples were spun down for 5 min at 13,000 xg at 4 EC and the supernatant was transferred to a new tube.

#### Complex formation of CAF-1-H3-H4 and PCNA without DNA

Histones H3-H4 (C110A,T71C) tetramers, labeled with AlexaFluor 488, were concentrated in 20 mM Hepes pH 7.5, 2 M NaCl, 1 mM EDTA, 1 mM TCEP to a final concentration of 79.4 μM using an Amicon Ultra-0.5 centrifugal concentrator with a molecular weight cut off of 10 kDa. CAF-1 WT or mutants were diluted to a concentration of 27.1 μM in 20 mM Hepes pH 7.5, 200 mM NaCl, 1 mM EDTA, 1 mM TCEP. CAF-1 was then mixed with the histones in a volumetric ratio of 3:1 to obtain samples with final concentrations of 20 μM CAF-1 and 10 μM H3-H4 tetramers. The NaCl concentration in these samples was around 650 mM._CAF-1–H3-H4 samples were mixed in order with buffer containing 20 mM Hepes pH 7.5, 60 mM NaCl, 1 mM TCEP and then PCNA (labeled with AlexaFluor 546, 185 μM stock) to obtain final concentrations of 1.5 μM CAF-1–H3-H4, 5.55 uM PCNA with a total of about 105 mM NaCl. Samples were diluted 2-fold in buffer containing 20 mM Hepes pH 7.5, 100 mM NaCl, 1 mM TCEP, 0.02% IGEPAL CA-630, and incubated at room temperature for 10 min. Samples were subjected to chemical crosslinking by addition of a final concentration of 0.2% glutaraldehyde (from a 2.5% stock in water). The samples were incubated at room temperature for 20 min before quenching the crosslinker by addition of 100 mM Tris pH 7.5 (from 1 M stock). Samples were spun down for 15 min at 13,000 xg at 4 °C and the supernatant was transferred to a new tube.

#### Crosslinking of CAF-1-PCNA on DNA at limiting PCNA concentrations

AlexaFluor546-labeled PCNAK164C (50 nM) was loaded onto nicked (with Nt.BspQ1) pUC19 plasmids (15 nM) by RFC (15 nM). The reaction was conducted at 30 °C for 5 minutes in buffer containing 20 mM HEPES-KOH pH 7.6, 130 mM NaCl, 0.01% IGEPAL CA-630, 1 mM TCEP, 10 mM MgCl2, 1 mM ATP. Then, CAF-1 or buffer control was titrated between 0-1 μM. The total NaCl concentration during the loading reaction and after addition of CAF-1, taking into account the contributions from each component, ranged between 100 and 110 mM. After 10 min at room temperature, the samples were diluted 4.5-fold by adding buffer containing 20 mM HEPES-KOH pH 7.6, 100 mM NaCl, 0.01% IGEPAL CA-630, 1 mM TCEP, before cross-linking with 0.2% glutaraldehyde. The cross-linking reaction took place at room temperature for 20 minutes, after which, it was quenched with a final concentration of 100 mM Tris-HCl pH 7.5. DNA was digested using Pierce™ Universal Nuclease for Cell Lysis (Thermo Fisher Scientific) diluted to 1:20 in 20 mM HEPES-KOH pH 7.6, 100 mM NaCl, 5 mM MgCl_2_, 1 mM TCEP and added to 10% of the crosslinking reaction volume. The digestion was quenched with 50 mM EDTA and immediately spun down for 5 minutes at 13,000 xg and 4°C to remove precipitates.

#### SDS-PAGE analysis of crosslinked samples

Crosslinked samples were mixed with 4x XT sample buffer and 20x XT reducing agent in appropriate volumetric ratios. These samples were loaded on 12% Criterion XT Bis-Tris gels in XT MOPS buffer. Gels were run at 20 mA until the samples have completely entered the gel and then at 40 mA until the gel run was complete (typically between 2 and 3 hours). Gels were run at room temperature, and additionally in the dark if components contained fluorophores. Gels were scanned for histones H3-H4 and/or PCNA fluorescence (depending on the assay) on an AMERSHAM ImageQuant 800 imager (Cytiva). Band intensity was calculated using the ROI manager tool in Image J/Fiji and plotted using GraphPad Prism. Where applicable, gels were subsequently stained with Coomassie blue and scanned on AMERSHAM ImageQuant 800 imager (Cytiva).

### Mass Photometry

Samples were prepared using crosslinking at stoichiometric conditions, the reactions (±1.2 mL final volume after EDTA quenching) were concentrated to 500 μL and loaded on a pre-equilibrated Superose 6 10/300 GL (Cytiva) column in buffer 20 mM HEPES-NaOH pH 7.5, 200 mM NaCl, 1 mM TCEP. Fractions were analyzed on SDS PAGE, the ones containing the complex of interest (Peak1 or Peak2) were pooled and concentrated to about 40 μL (Abs280 close to 0.5). The samples were diluted 10 to 20-fold in 20 mM HEPES-NaOH pH 7.5, 100 mM NaCl, 1 mM TCEP right before measuring on a Refeyn OneMP instrument (Refeyn Ltd.). For each measurement, 13 μL of this buffer was first placed into the CultureWell gaskets wells (Grace Biolabs) placed into the Microscope coverslips (24 mm × 50 mm; Paul Marienfeld GmbH). After adjusting the focus, 2 μL of sample was mixed in. Movies were recorded for 60 seconds at 100 frames per second. A calibration measurement under the same conditions was performed roughly every 15 measurements using an in-house prepared protein standard mixture: IgG4Δhinge-L368A (73 kDa), IgG1-Campath (149 kDa), apoferritin (479 kDa), and GroEL (800 kDa). Data were processed using DiscoverMP (Refeyn Ltd.) with bin width adjusted to 10, and each sample retrieved about 1500-3000 counts. Figures were prepared with the Refeyn instrument and edited in Illustrator.

### Fluorescence polarization

Fluorescence Polarization assays were carried out in 25 mM TRIS pH 7.5, 300 mM NaCl, 5% glycerol, 1 mM EDTA, 0.01% NP-40 (added fresh), 0.01% CHAPS (added fresh), 1 mM DTT (added fresh). Binding reactions were prepared by mixing 10 nM of Alexa488-labeled H3-H4 (H3 C110A-H4 E63C) and increasing amounts of CAF-1, Polε, Polδ or RPA in a final volume of 30 μL in CORNING low flange 384 well black microplates (CLS3575). Binding data were measured using a CLARIOStar (BMG LabTech) plate reader. The data was analyzed and plotted using Microsoft Excel and GraphPad Prism. Kd-values were calculated using a one site binding curve in GraphPad Prism. Representative curves are shown from one experiment (three independent measurements) and were repeated at least two times in triplicates.

### NAQ assay

This refers to Supplementary Figure S5B. The nucleosome assembly reaction was carried out at 200 nM of 207 bp DNA, 200 nM xenopus octamer maleimide AlexaFluor-647 (AF647) labeled on H2B T112C (containing H3 C110A mutant) and 500 nM CAF-1, Polε or RPA. After the assembly reaction, the samples were diluted to a DNA concentration of 50 nM in 100 μl digestion reactions. 25U of MNase enzyme was added in a final buffer containing 50 mM TRIS pH 7.9, 5 mM CaCl_2_. After incubation at 37°C for 10 min, the reactions were quenched with 10 μl of 500 mM EDTA, pH 8. The DNA was then purified using a modified protocol of the MinElute kit from QIAGEN. 550 μl of PB buffer and 10 μl of 3 M sodium acetate were added to each sample and they were incubated at room temperature for 10 min. At this point, 50 ng of DNA loading control (or reference band, a 621 bp DNA fragment) was added to each tube. The samples were applied to the MinElute spin column and washed as prescribed by QIAGEN. The DNA was eluted with 10 μl of water. 2.5 μl were loaded on a 6% PAGE gel. The gel was run for 45 min at 200 V in 0.5x TBE buffer at room temperature. Gels were stained with SybrGOLD for DNA and imaged on an AMERSHAM ImageQuant 800 (Cytiva).

### Cell culture, genome editing and western blot

Mouse ESCs used in this study were derived from the E14JU cell line with a 129/Ola background. For genome editing and next-generation sequencing experiments, ESCs were grown on gelatin-coated dishes (0.2 %) in serum+LIF conditions at 37 °C with 5 % CO2. Media was prepared by supplying DMEM-GlutaMAX-pyruvate with fetal bovine serum (15 %), LIF (made in house), 1x non-essential amino acids (Gibco), 1x penicillin/streptomycin (Gibco) and 2-beta-ME (0.1 μM). Cells were passaged using Trypsin-EDTA (Gibco) or TrypLE (Gibco). Cells were routinely tested for mycoplasma contamination. For genome editing *Chaf1a*-dTAG cells were generated by CRISPR-Cas9 using the SpCas9(BB)-2A-Puro (PX459) V2.0 plasmid (addgene #62988) as described in (64) with sgRNA#1 (Supplementary Table S5), which target the Chaf1a gene at the beginning of the ORF and a Chaf1-linker-dTAG homology donor plasmid. Cells were transfected using Lipofectamine 3000 reagent (Invitrogen) using 0.5 μg of sgRNA-plasmid and 2 μg of donor plasmid. Cells were sparsely seeded on a 10 cm dish 24 hours posttransfection and selected with Puromycin (2 μg/mL) for 48 h. Thereafter, cells were expanded and genotyped with primers #1 and #2 (Supplementary Table S5). Positive clones were analyzed by Sanger sequencing with primers #3 and #4 (Integrated DNA Technologies, Supplementary Table S5) and degradation upon dTAG-13 (Tocris, 6605) treatment was confirmed by Western Blot by a-Chaf1a antibody (65). Fractionation cell extracts were prepared as in (66). Western Blotting was performed as described in (67).

### Immunofluorescence

Cells treated with DMSO or dTAG-13 for 4 hours,were pulsed in EdU-containing media (10 μM) for 10 min and immediately fixed for 15 min in 4% PFA at RT and stored in PBST (PBS with 0.3% Triton X-100). Primary antibody H4K20me0 was added at the concentration of 1:1000 in PBST with 5% donkey serum and incubated overnight. Incubation was followed by 3 washes in PBS and secondary antibody was then added in PBST. Samples were incubated with the secondary antibody in the dark at RT for 1h. After 3 washes, samples were stained with DAPI (1:10000) in PBST. Images were acquired with a ScanR high-content screening microscope (Olympus). Automated and unbiased image analysis was carried out with the ScanR analysis software (version 2.8.1). Individual cells were identified based on DAPI staining and mean pixel intensity was measured for each channel. Data were exported and processed using Spotfire software (version 10.5.0; Tibco). Statistical analysis and visualization of results was done using using R (v4.1.2) in RStudio (v2021.9.2.382).

### SCAR-seq

A step-by-step protocol is available (68). Briefly, nascent SCAR-seq samples were prepared from *Chaf1a*-dTAG cells in three biological replicates for each histone PTM. Cells treated with DMSO or dTAG-13 for 2 hours, were pulsed in EdU-containing media (10 μM) for 30 min and harvested immediately. For sample collection, media was aspirated, plates washed 2x with RT PBS and ice-cold PBS was added to the dishes. Cells were scraped in a cold room and collected by centrifugation, followed by nuclei isolation. Nuclei were aliquoted, snap-frozen and stored at −80° C until further use. For MNase digest, nuclei were counted manually using Kova Glasstic Slides and 2 U MNase (Worthington) were added per 1×106 nuclei. Digests were performed at 30° C for 20 minutes. For native ChIP, 30-50 μg of chromatin was used per sample and incubated with antibodies in a total volume of 600 μL overnight at 4° C with H3K27me3 antibody (Cell Signaling, 9733) or H4K20me0 antibody (Abcam, ab227804). Magnetic beads (anti-rabbit IgG Dynabeads, invitrogen) were added the next morning and samples were incubated for 2 hours. After three washes each with ice-cold RIPA buffer and RIPA 0.5M NaCl buffer, DNA was eluted and purified using the MinElute Reaction Cleanup kit (Qiagen). Mononucleosomal-sized fragments were isolated by double sided size selection with AMPure XP beads (Beckman Coulter). EdU-labelled DNA fragments were biotinylated using Click-iT chemistry as reported above but using Biotin-TEG-Azide (Berry & Associates) instead of Picolyl-azide-PEG4-Biotin. Libraries were prepared using the KAPA Hyper Prep Kit (Roche). Biotinylated fragments were captured using Dynabeads MyOne Streptavidin (invitrogen) and EdU strands were purified by performing NaOH washes. Libraries were amplified in 9-11 PCR cycles. Libraries with mononucleosomal-sized inserts were isolated by double-sided size selection with AMPure XP beads (Beckman Coulter), followed by a second clean-up with 1.0x AMPure XP beads. Fragment distribution of libraries was checked on a a Fragment Analyzer system (Agilent). Stranded input samples were prepared in parallel with SCAR-seq samples. Samples were sequenced single end (75bp) on a NextSeq500 instrument (Illumina).

Reads were processed, mapped and histone partition signal was computed as described previously(68). Briefly, for each strand the SCAR normalized signal (CPM) was computed in 1kb bins and smoothed in a uniform blur considering the neighbouring 30 bins on each side. For each 1kb window, the signal from its corresponding SCAR input was subtracted and negative values were set to zero. Input corrected windows with CPM < 0.3 on both strands were filtered out and not considered for further analyses. The final partition score for each 1kb window was calculated as: Partition = (F - R)/(F + R) where F and R correspond to the number of normalized and input-corrected reads for the forward and reverse strand, respectively. The partition value relates to the ratio of histones with a specific modification being segregated to the nascent forward (Partition > 0) or nascent reverse (Partition < 0) strand within each window respectively. Okazaki-seq replication fork directionality (RFD) scores and filtered initiation zones (IZs) for mESC were taken from (67) and used to define replication via leading or lagging strand mechanism. The RFD score in Okazaki-seq is calculated like SCAR-seq partition scores but subtracting the forward (F) strand signal from the reverse (R) strand signal instead: RFD = (R - F)/(F + R).

The average partition signal from replicate 1 was used for visualization purposes in Partition line plots (Figure 6). To visualize the total reads in SCAR-seq, total mm10 mouse read counts were spike-in normalized to dm6 drosophila read counts as described in (69) By using the uniquely mapping, deduplicated reads in millions, the EdU-enriched Input samples (“ClickedInputs”) was used as reference for relative spikeIn abundance and EdU labelling efficiency. To visualize global signal in SCAR-seq, number of uniquely mapped, deduplicated mm10 reads of the SCAR sample (in million reads) were normalized to DMSO for each mark and replicate and plotted in replicates using R (v4.1.2) in RStudio (v2021.9.2.382).

### End-point DNA replication with yeast replisome

These were carried out as in (9), all stock protein concentrations were determined by Bradford analysis. MCM was loaded onto 5.8 Kb ARS1 plasmid in 30 μL reaction volumes, to final concentrations of 22.5 nM ORC, 100 nM Mcm2/7-cdt1, 45 nM Cdc6 and 4 nM plasmid DNA template, in buffer containing 25 mM Hepes-KOH pH 7.6, 100 mM potassium glutamate, 10 mM magnesium acetate, 0.02% IGEPAL CA-630, 5% glycerol, 5 mM ATP, 0.1 mg/mL BSA, 1 mM DTT. This reaction was incubated at 30°C for 20 minutes. After origin licensing, DDK was added to 25 nM and further incubated at 30°C for 30 minutes. The replication reaction was initiated by addition of FF500 buffer (50 mM HEPES-KOH pH 7.6, 500 mM potassium glutamate, 20 mM magnesium acetate, 0.02% IGEPAL CA-630, 2 mM DTT, 6 mM ATP, 0.2 mg/mL BSA, 0.4 mM CTP, GTP, UTP each, 0.16 mM dGTP, dATP, dTTP, dCTP and 40 nM α32P-dCTP), followed by replication proteins in a master mix added to final reaction concentrations of 30 nM Dpb11, 40 nM cdc45, 210 nM GINS, 20 nM S-CDK, 5 nM Mcm10, 25 nM Sld3/7, 50 nM Sld2, 20 nM Polε, 100 nM RPA, 20 nM Polα, 20 nM Ctf4, 20 nM TopoII, and another protein master mix added to final reaction concentrations of 20 nM Mrc1, 20 nM Csm3/Tof1, 10 nM TopoI, 20 nM RFC, 20 nM PCNA, 10 nM Polδ. The replication reaction was conducted at 30 °C for 40 minutes. After this, the reaction was quenched by addition of 50 mM EDTA to 2x dilution. Samples were cleaned-up for unincorporated nucleotides using MicroSpin G-50 columns (Cytiva), after which they were denatured in 100 mM NaOH, 2% sucrose, bromocresol green as loading dye and 15 μL samples were run on 0.7% alkaline agarose gels for 18 hours at 45 V. The next day, DNA was precipitated on gel by treatment with ice cold 5% TCA for 2 cycles of 15 minutes with TCA refreshment. The gel was dried in 2x chromatography Whatman paper and towel paper sandwich with a weight on top for 30 minutes, to remove excess moisture. After that, the Whatman paper gel sandwich was moved to a gel dryer for 2.5 hour at 55 °C. Gel was exposed to a phosphor screen for 2 days using Amersham Typhoon Biomolecular Image.

### Pulse-chase experiments

These were carried out as in (9), all stock protein concentrations were determined by Bradford analysis. MCM was loaded onto 5.8 Kb ARS1 plasmid in 150-300 μL reaction volumes, to final concentrations of 22.5 nM ORC, 100 nM Mcm2/7-cdt1, 45 nM Cdc6 and 4 nM plasmid DNA template, in buffer containing 25 mM Hepes-KOH pH 7.6, 100 mM potassium glutamate, 10 nm magnesium acetate, 0.01% IGEPAL CA-630, 5 mM ATP, 0.1 mg/mL BSA, 1 mM DTT. This reaction was incubated at 30°C for 30 minutes. After origin licensing, DDK was added to 25 nM and further incubated at 30°C for 30 minutes. Replication proteins in a master mix were added to final pulse reaction concentrations of 30 nM Dpb11, 40 nM cdc45, 210 nM GINS, 20 nM S-CDK, 5 nM Mcm10, 25 nM Sld3/7, 50 nM Sld2, 20 nM PolE, 100 nM RPA, 40 nM PolA, 20 nM Ctf4, followed by another protein master mix added to final pulse reaction concentrations of 20 nM Mrc1, 20 nM Csm3/Tof1, 10 nM TopoI. This reaction was then split into the pulse mixes containing 20 nM RFC, 20 nM PCNA, 180 nM CAF-1 (or corresponding storage buffers for control reactions), FF500 pulse buffer (50 mM HEPES-KOH pH 7.6, 500 mM potassium glutamate, 20 mM magnesium acetate, 0.02% IGEPAL CA-630, 2 mM DTT, 6 mM ATP, 0.2 mg/mL BSA, 0.4 mM CTP, GTP, UTP each, 0.16 mM dGTP, dATP, dTTP, 4 μM dCTP and 66 nM α32P-dCTP. Pulse reaction was conducted at 30 °C for 3 min 20 sec, when the chase (0.6 mM dCTP, dGTP, dATP, dTTP) was added. Time points were taken (15 μL) at 4, 5, 6 and 7 min and replication reaction was quenched by addition to 2x dilution in 50 mM EDTA. Samples were cleaned-up for unincorporated nucleotides using MicroSpin G-50 columns (Cytiva), after which they were denatured in 10 mM NaOH, 2% sucrose, bromocresol green as loading dye and 14 μL samples were run on 0.7% alkaline agarose gels for 18 hours at 45 V. The next day, DNA was precipitated on gel by treatment with ice cold 5% TCA for 2 cycles of 15 minutes with TCA refreshment. The gel was dried in 2x chromatography Whatman paper and towel paper sandwich with a weight on top for 30 minutes, to remove excess moisture. After that, the Whatman paper gel sandwich was moved to a gel dryer for 2.5 hour at 55 °C. Gel was exposed to a phosphor screen for 2 days using Amersham Typhoon Biomolecular Image. Data analysis was performed using ImageQuant TL software. Product length was determined with respect to ladder product sizes of Lambda x HindIII DNA (Promega). The front (fastest migrating population) was identified when manually assigning the peaks on ImageQuant software. Data were fit using linear regressions, and the slope determined replication rates in GraphPad prism.

## AUTHORS CONTRIBUTIONS

Conceptualization: C.R., B.V.E., L.K., A.G., F.M.

Data Curation: C.R., B.V.E., L.K., F.G., F.M.

Formal Analysis: C.R., B.V.E., L.K., F.G., A.E.E.V., I.R., P.H.L.K., F.M.

Funding acquisition: A.G., F.M.

Investigation: C.R., B.V.E., L.K., F.G., A.E.E.V., I.R., G.R., P.A., F.M.

Methodology: C.R., B.V.E., L.K., F.G., A.E.E.V., I.R., A.B., T.v.L., P.A., F.M.

Project administration: F.M.

Software: P.H.L.K.

Supervision: R.A.S., W.d.L., N.H.D., A.G., F.M.

Validation: C.D.v.F., G.L.

Visualization: C.R., B.V.E., L.K., F.G., F.M.

Writing – original draft: C.R., B.V.E., F.M.

Writing – Review & Editing: all authors

## ACKNOLEDGMENTS

We thank John Diffley, Peter Burgers, Xiaodong Zhang and Rafael Fernández Leiro for sharing reagents, Titia Sixma and Shreya Dharadhar for help with RFCΔN purification, Puck Knipscheer and Juan Garaycoechea for discussions and critical reading of the manuscript, Anke Sparmann at Life Science Editors for editing services (www.lifescienceeditors.com), Jamie Barnett for designing and sharing primers for NGS, Jan Dreyer for discussions on the PCNA-NAQ assay, and The Histone Source at Colorado State University for lyophilized histones.

