## Supplemental Information for "CAF-1 deposits newly synthesized histones during DNA replication using distinct mechanisms on the leading and lagging strands"

**This PDF file includes:**

Supplementary Figures S1 to S7

Supplementary Tables S1 to S5

### Supplementary Figures

Supplemental Figure S1

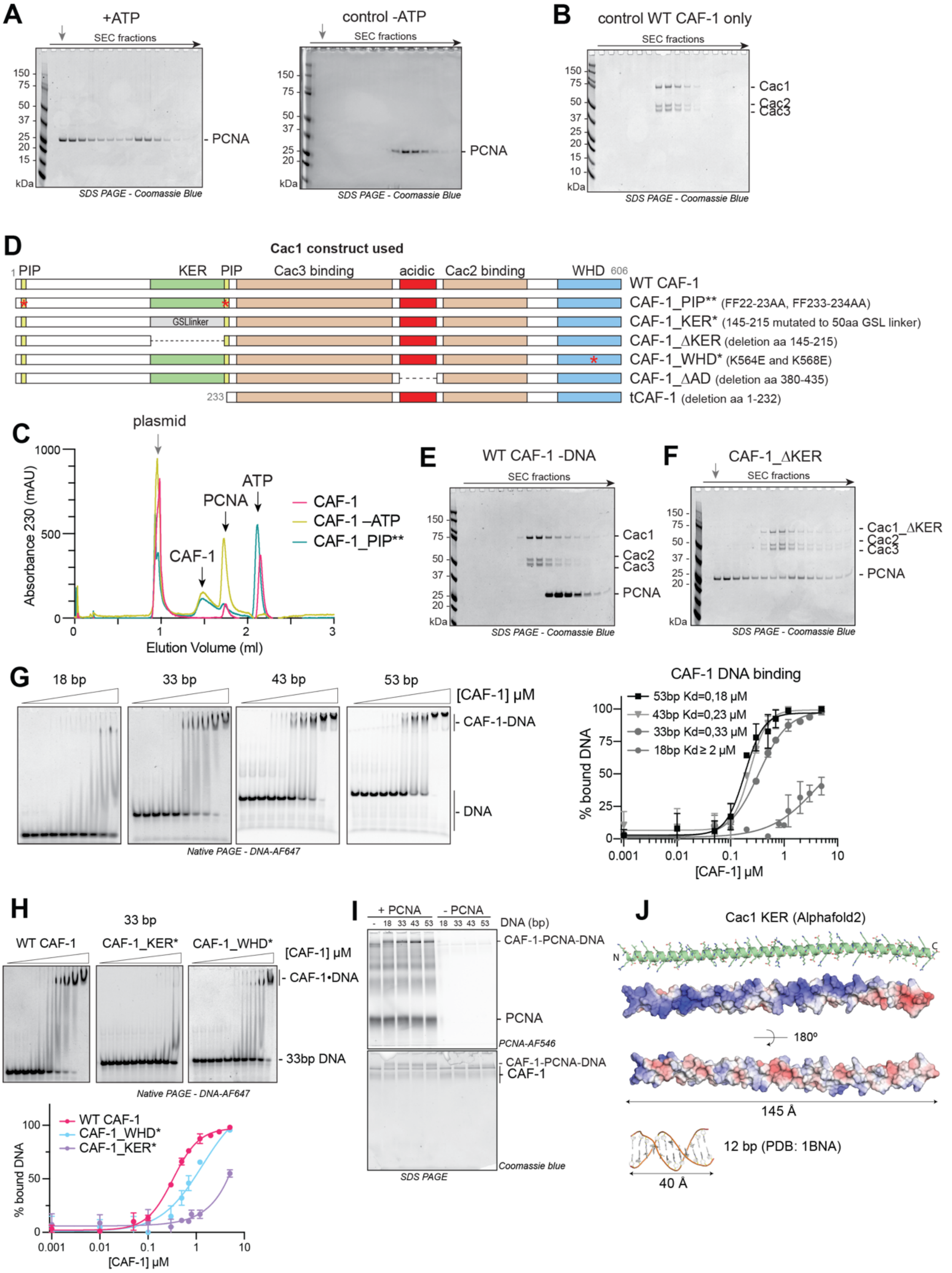


**Supplementary Figure S1**

**A)** SDS PAGE following separation on SEC of a PCNA loading reaction on DNA plasmids (on the left) and the control omitting ATP (on the right). The grey arrow indicates the elution volume of the plasmid DNA. **B)** SDS PAGE following separation on SEC of WT CAF-1 only. **C)** Chromatogram (230 nm signal) of the Superose 6 runs whose SDS PAGE gels are shown in Figure 1A-C. The grey arrow indicates the elution volume of the plasmid DNA.

Elution volume for CAF-1, PCNA and ATP are marked by black arrows. **D)** Cartoons of the Cac1 (large CAF-1 subunit) domains and the mutants used in this study. All complexes contain Cac2 and Cac3, in addition to the indicated Cac1 construct. **E)** SDS PAGE following separation on SEC of a control reaction where the DNA plasmid was omitted, from a CAF-1-PCNA binding experiment as in Figure 1A. **F)** SDS PAGE following separation on SEC of a CAF-1-PCNA binding reaction on nicked DNA plasmid using a CAF-1_ΔKER mutant. The grey arrow indicates the elution volume of the plasmid DNA. **G)** EMSA experiments and quantification of CAF-1 binding to double-stranded DNA fragments. Each 18-33-43-53 bp DNA fragment carries a AF647 fluorophore for detection and quantification. Disappearance of the unbound DNA band was used to calculate binding at different CAF-1 concentrations. Binding affinities Kd result from fitting the calculate data in GraphPad. 18 bp data is fitted with a One Site total binding curve, while the data with 33-43-53 bp DNA was fit accounting for cooperativity. The Hill coefficients obtained for these curves are 1.6, 2.7, 2.4 respectively. Means ±SD is shown for each data point. At least three replicates were done for each experiment. **H)** EMSA experiments and quantification of binding to a 33-bp double-stranded DNA fragment of CAF-1_KER* and CAF-1_WHD* mutants. Disappearance of the unbound DNA band was used to calculate binding at different CAF-1 concentrations. Means ±SD is shown for each data point. At least three replicates were done for each experiment. **I)** Crosslinking experiment between CAF-1 (3 $\mu$M) and labeled PCNA (4.5 $\mu$M) on DNA fragments (1.5 $\mu$M) of various sizes. DNA was not digested in these reactions. RFC and ATP were not added to actively load PCNA. These are the full gels of Figure 1G. **J)** Alphafold model of the KER domain in Cac1 (residues 128-226). Cartoon with sticks and electrostatics are shown. The electrostatics are calculated with the APBS plugin in Pymol. Blue is +5kEV and Red is -5kEV. A 12bp structure of B-DNA (PDB: 1BNA) is shown at the same scale of the KER domain for length comparison.

### Supplemental Figure S2

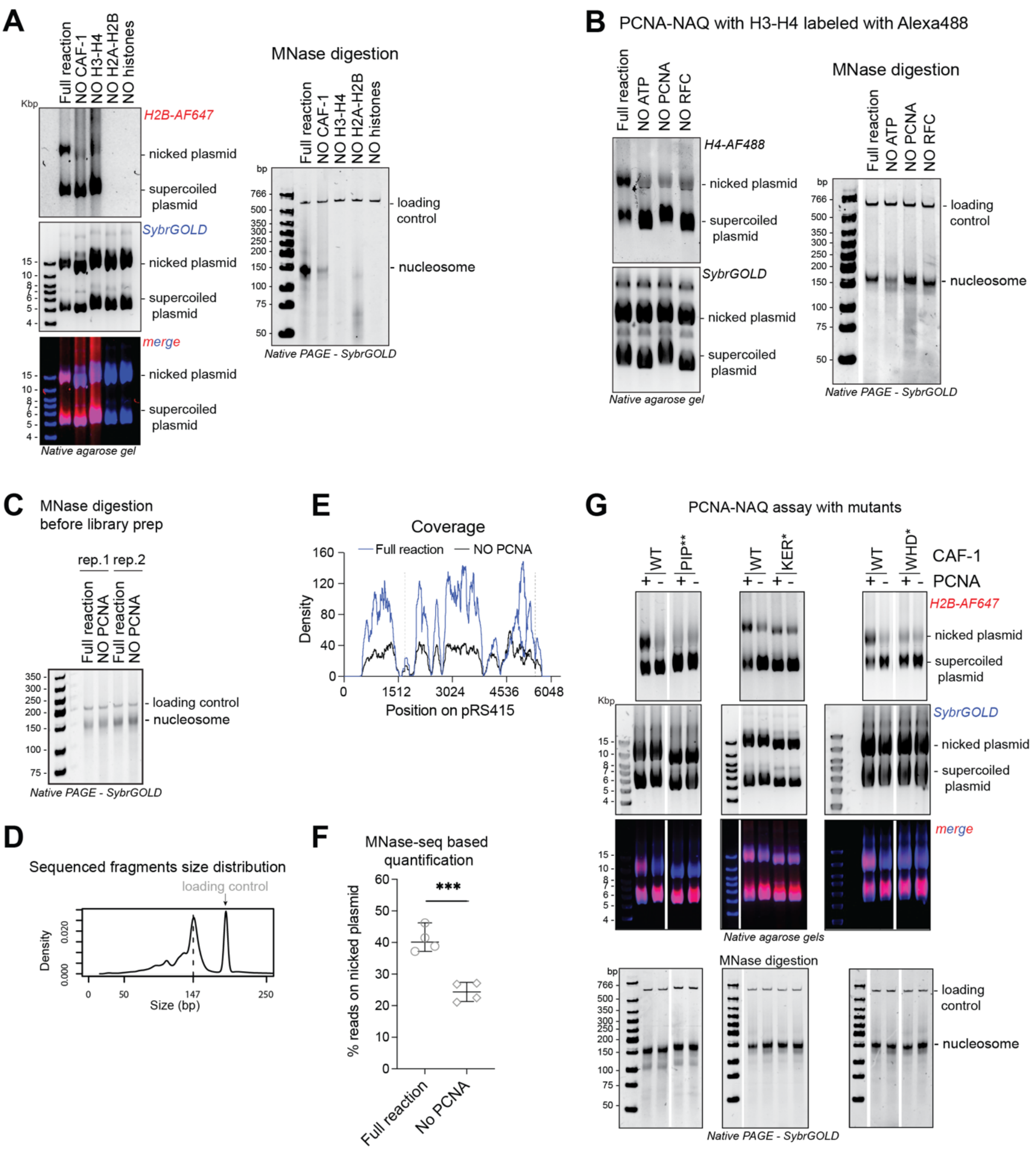


**Supplementary Figure S2**

**A)** Left panel: Native agarose gel of control PCNA-NAQ assay conditions. Fluorescence signal for H2B-T112C labeled with AF647 (H2B-AF647) or DNA (SybrGOLD), and their overlay are shown. H2B fluorescence on the nicked plasmid (top panel) represents PCNA-dependent histone deposition. Right panel: Native PAGE stained with SybrGOLD of protected DNA fragments following MNase digestion. 150bp DNA fragments are characteristic of nucleosomal DNA, a 621bp loading control is used to monitor DNA retrieval during the purification procedure.
**B)** Left panel: Native agarose gel of control PCNA-NAQ assay conditions where instead of fluorescently labeled H2A-H2B, we used labeled H3-H4. Fluorescence signal for H4-E63C labeled with AF488 (H4-AF488) is shown. Subsequently we stained with SybrGOLD to image DNA (SybrGOLD also emits at 488 nm). H4 fluorescence on the nicked plasmid (top panel) represent PCNA-dependent histone deposition. Right panel: Native PAGE stained with SybrGOLD of protected DNA fragments following MNase digestion. 150bp DNA fragments are characteristic of nucleosomal DNA, a 621bp loading control is used to monitor DNA retrieval during the purification procedure.
**C)** MNAse digestion gel of PCNA-NAQ assay samples obtained for NGS analysis. These reactions contained a nicked and supercoiled plasmid with different sequences (pRS415 or pLox3). We used a 207 bp DNA loading

control containing a 601-widom sequence. **D)** Fragment size distribution of the sequenced reads confirms the dominance of a 150 bp size after MNase digestion and sequencing. **E)** Example of reads coverage on the nicked plasmid when pRS415 was used in the full reaction or in a negative control reaction omitting PCNA. Dashed lines show the sites of nicking. **F)** Quantification of the PCNA-dependent nucleosome assembly activity based on the NGS reads of WT CAF-1 and a no-PCNA control reaction. The percentage of reads on the nicked plasmid is shown over the total number of reads (for both plasmids). Means ±SD is shown, and an unpaired t-test was applied to determine statistical significance, *** p>0,001. **G)** Native agarose gel (top) of control PCNA-NAQ assay conditions with CAF-1 mutants. Fluorescence signal for H2B-T112C labeled with AF647 (H2B-AF647) or DNA (SybrGOLD), and their overlay are shown. H2B fluorescence on the nicked plasmid (top panel) represents PCNA-dependent histone deposition. Bottom: Native PAGE stained with SybrGOLD of protected DNA fragments following MNase digestion. 150bp DNA fragments are characteristic of nucleosomal DNA, a 621bp loading control is used to monitor DNA retrieval during the purification procedure. Quantifications of H2B fluorescence are shown in Figure 2C.

Supplemental Figure S3

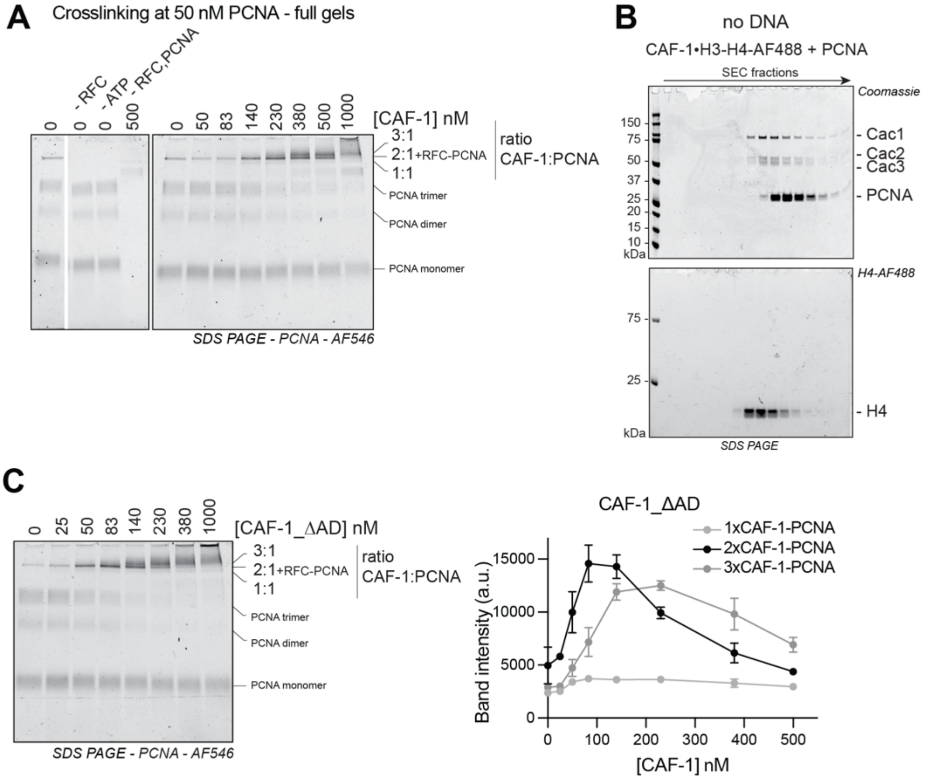


**Supplementary Figure S3**

**A)** Full gels of data in Figure 3C. PCNA fluorescence scan of SDS-PAGE of crosslinking experiment after DNA digestion, of reactions containing 50 nM PCNA, 15 nM RFC, 15 nM pUC19 and increasing CAF-1 concentrations. **B)** SDS PAGE after SEC of a reaction containing PCNA (30 μM), and CAF-1 preloaded with labeled H3-H4 (5 μM). No DNA or RFC is present in these reactions. After SDS PAGE run, we scan the gel for the H4-488 nM fluorescence (bottom), followed by staining with Coomassie (top). **C)** PCNA fluorescence scan of SDS-PAGE following protein-protein crosslinking after DNA digestion. These reactions contain 50 nM PCNA, 15 nM RFC, 15 nM pUC19 and increasing CAF-1_ΔAD concentrations. On the right: quantification of the CAF-1-PCNA bands. Mean ± SD is shown of three replicates.

Supplemental Figure S4

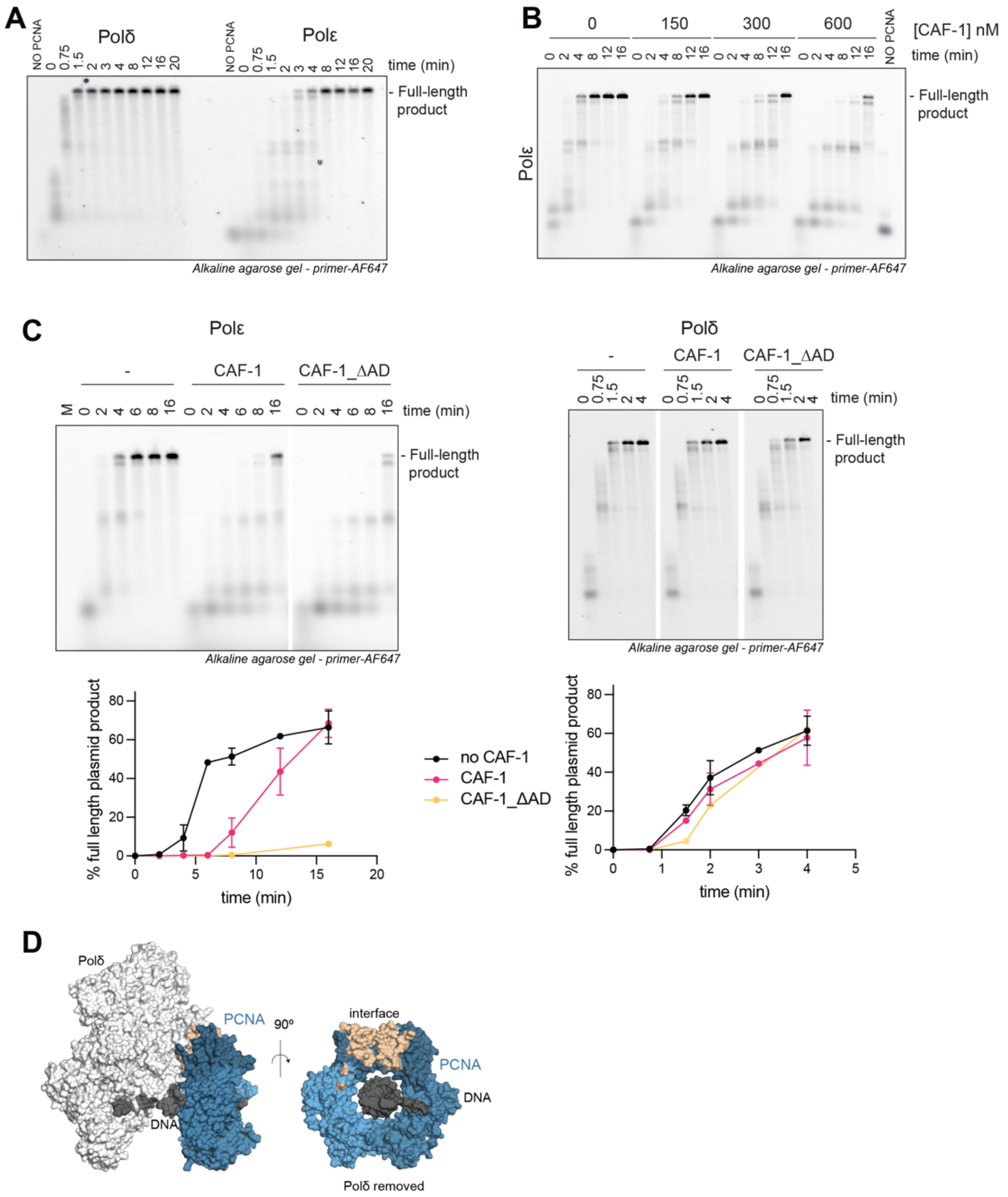


**Supplementary Figure S4**

**A)** Fluorescence (primer signal) scan of denaturing alkaline agarose gel of primer extension reactions with Polδ or Polε. The polymerases were at 120 nM, PCNA 480 nM. Both polymerases are active with different kinetics. Both polymerases depend on PCNA for activity. The NO-PCNA control lanes contain reactions that incubated for 20 minutes. **B)** Fluorescence (primer signal) scan of denaturing alkaline agarose gel of primer extension reactions with Polε in the presence of increasing amounts of WT CAF-1 (150-300-600 nM). **C)** Fluorescence (primer signal) scan of denaturing alkaline agarose gel of primer extension reactions with Polε (left) or Polδ (right) in the presence of CAF-1 WT or CAF-1_ΔAD (300 nM). Bottom: Quantification of the full-length product band relative to the total fluorescence in each lane (expressed as percentages). Mean ±SD are shown for three replicates. **D)** Surface visualization of Polδ (in white) bound to PCNA (blue) on DNA (dark gray) from PDB 7KC0. In wheat we show the PCNA residues involved in the Polδ interaction. On the right panel Polδ is not displayed.

Supplemental Figure S5

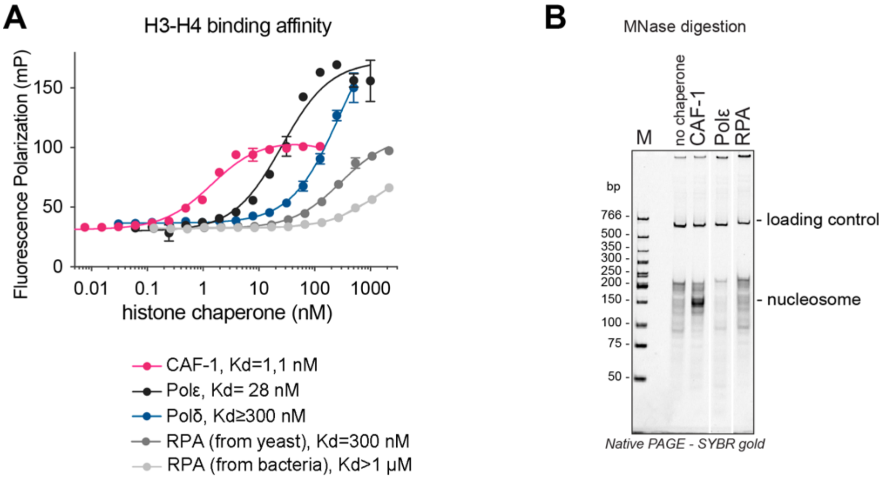


**Supplementary Figure S5**

**A)** Fluorescence polarization experiment to test binding to H3-H4 (where H4 is labeled at E63C using AF488). CAF-1, Polε, Polδ, RPA expressed in yeast (used in Figure 7A) and RPA expressed in bacteria (used in Figure 4-5) were titrated to a solution containing 10 nM of labeled H3-H4. The data points were fit to a one site binding curve and the Kd were calculated in GraphPad. **B)** MNase digestion of a NAQ reaction where each histone chaperone was incubated with the histone octamer and subsequently with 207 bp DNA. Bands at 150 bp represent nucleosomes and these are observed only in the presence of CAF-1. RPA produced from yeast cells was used. A 621 bp loading control DNA is used to control for sample retrieval during the DNA purification step.

Supplemental Figure S6

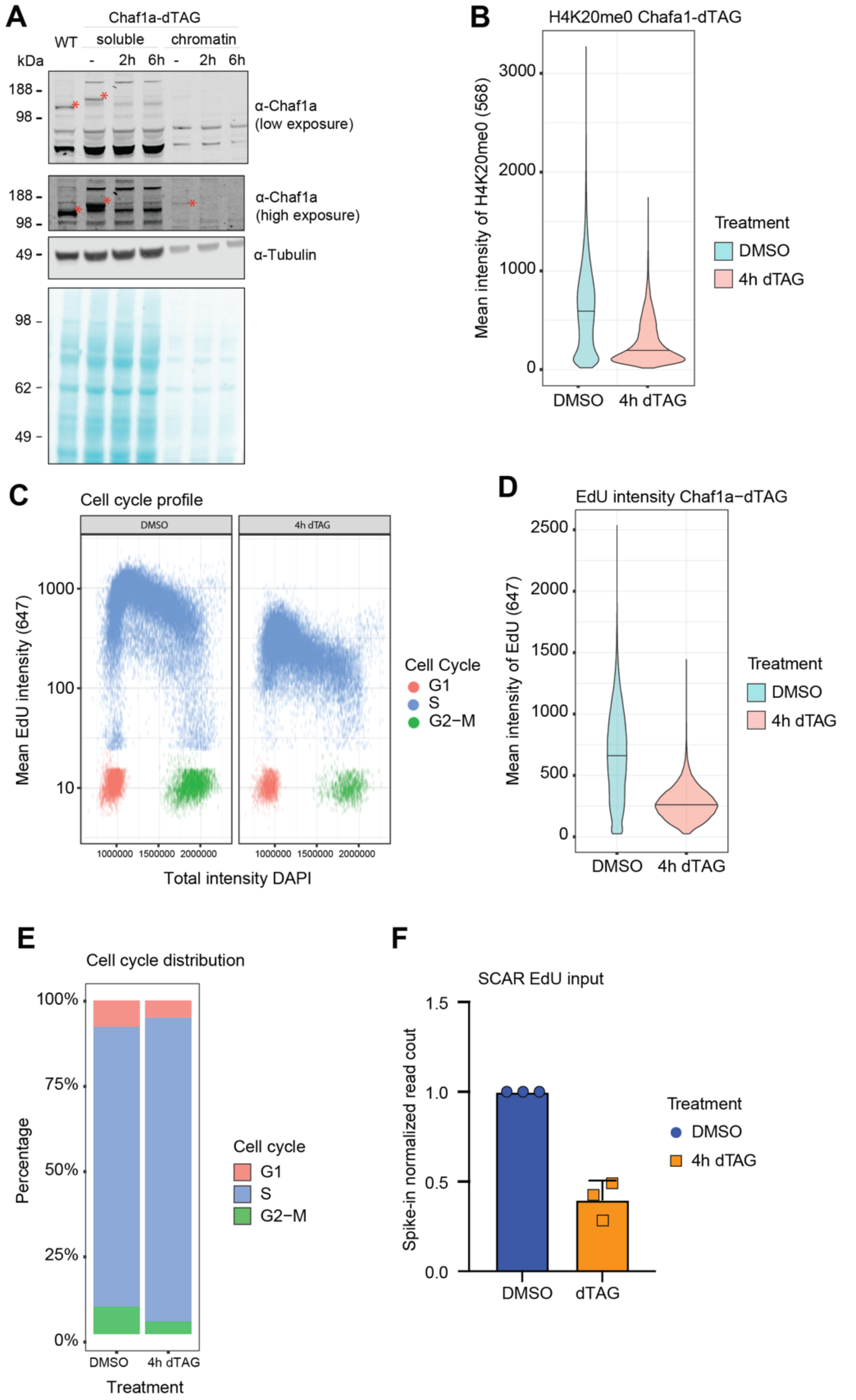


**Supplementary Figure S6**

**A)** Western blot analysis of soluble and chromatin fractions of mES cells treated with DMSO and dTAG for the indicated times. A WT cell line is shown as a control. **B)** Immunofluorescence results of mean H4K20me0 intensities in mESC upon CAF-1 depletion with dTAG shows a decrease in H4K20me0. **C)** Immunofluorescence results of mean EdU intensity vs total DAPI intensity. **D)** Mean EdU intensity in DMSO vs dTAG treated cells. **E)** Cell cycle distribution based on mean EdU intensity and total EdU intensity. Immunofluorescent data are represented from two replicates. **F)** Spike-in normalized input reads shows decreased EdU incorporation after depletion of CAF-1.

Supplemental Figure S7

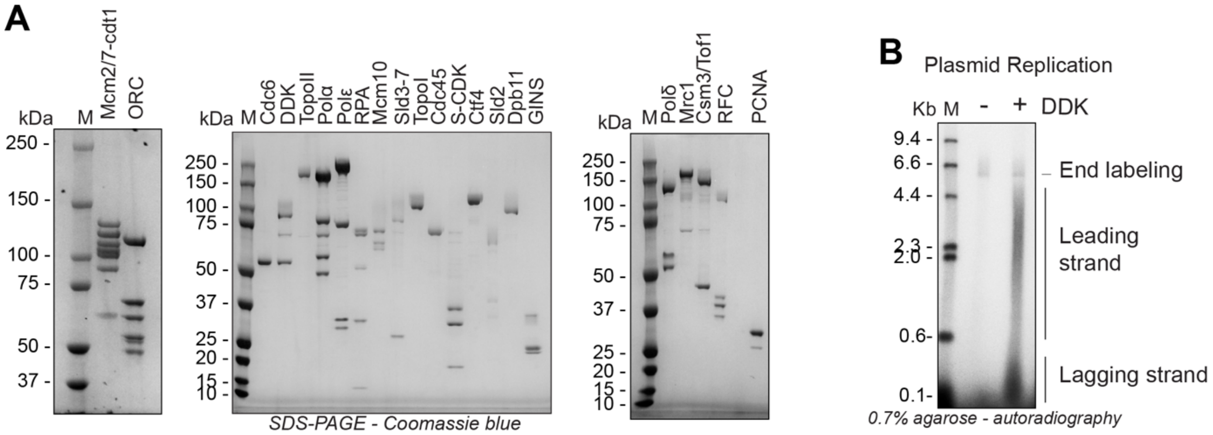


**Supplementary Figure S7**

**A)** SDS PAGE of protein preps that were used to reconstitute the yeast replisome. **B)** Autoradiography scan of denaturing agarose gel separation using DNA replication products, from an end-point plasmid replication experiment containing all yeast replisome components or, as control, omitting DDK.

### Supplementary Information

Supplementary Table S1: Primers used for EMSA.

| DNA primers | Reference |
|---|---|
| DNA oligo 18mer forward: GTCTACGAGCAATTGAGC | Mattiroli et al., 2017 |
| DNA oligo 18mer reverse: GCTCAATTGCTCGTAGAC | Mattiroli et al., 2017 |
| DNA oligo 33mer forward: GCTGTCTACGAGCAATTGAGCGGCCTCGGCACC | Mattiroli et al., 2017 |
| DNA oligo 33mer reverse: GGTGCCGAGGCCGCTCAATTGCTCGTAGACAGC | Mattiroli et al., 2017 |
| DNA oligo 43mer forward, 5' conjugated to AlexaFluor 647: CTAGAGCTGTCTACGAGCAATTGAGCGGCCTCGGCACCGGGAT | This paper |
| DNA oligo 43mer reverse: ATCCCGGTGCCGAGGCCGCTCAATTGCTCGTAGACAGCTCTAG | This paper |
| DNA oligo 53mer forward, 5' conjugated to AlexaFluor 647: CGGTGCTAGAGCTGTCTACGAGCAATTGAGCGGCCTCGGCACCGGGATTCTGA | This paper |
| DNA oligo 53mer reverse: TCAGAATCCCGGTGCCGAGGCCGCTCAATTGCTCGTAGACAGCTCTAGCACCG | This paper |

Supplementary Table S2, DNA sequences for plasmids and linear fragments used in MNase-seq

| Name | DNA sequence |
|---|---|
| pLox3 | AACGACCTACACCGAACTGAGATACCTACAGCGTGAGCTATGAGAAAGCGCCACGCTTCCCGAAGGG<br>AGAAAGGCGGACAGGTATCCGGTAAGCGGCAGGGTCGGAACAGGAGAGCGCACGAGGGAGCTTCC<br>AGGGGGAAACGCCTGGTATCTTTATAGTCCTGTCGGGTTTCGCCACCTCTGACTTGAGCGTCGATTTTT<br>GTGATGCTCGTCAGGGGGGCGGAGCCTATGGAAAAACGCCAGCAACGCGGCCTTTTTACGGTTCCTG<br>GCCTTTTGCTGGCCTTTTGCTCACATGTTCTTTCCTGCGTTATCCCCTGATTGACTTGGGTCGCTCTTCCT<br>GTGGATGCGCAGATGCCCTGCGTAAGCGGGTGTGGGCGGACAATAAAGTCTTAAACTGAACAAAATA<br>GATCTAAACTATGACAATAAAGTCTTAAACTAGACAGAATAGTTGTAAACTGAAATCAGTCCAGTTATG<br>CTGTGAAAAAGCATACTGGACTTTTGTTATGGCTAAAGCAAACTCTTCATTTTCTGAAGTGCAAATTGC<br>CCGTCGTATTAAAGAGGGGCGTGGCCAAGGGCATGTAAAGACTATATTCGCGGCGTTGTGACAATTTA<br>CCGAACAACTCCGCGGCCGGGAAGCCGATCTCGGCTTGAACGAATTGTTAGGTGGCGGTACTTGGGT<br>CGATATCAAAGTGCATCACTTCTTCCCGTATGCCCAACTTTGTATAGAGAGCCACTGCGGGATCGTCAC<br>CGTAATCTGCTTGCACGTAGATCACATAAGCACCAAGCGCGTTGGCCTCATGCTTGAGGAGATTGATG<br>AGCGCGGTGGCAATGCCCTGCCTCCGGTGCTCGCCGGAGACTGCGAGATCATAGATATAGATCTCACT<br>ACGCGGCTGCTCAAACTTGGGCAGAACGTAAGCCGCGAGAGCGCCAACAACCGCTTCTTGGTCGAAG<br>GCAGCAAGCGCGATGAATGTCTTACTACGGAGCAAGTTCCCGAGGTAATCGGAGTCCGGCTGATGTT<br>GGGAGTAGGTGGCTACGTCTCCGAACTCACGACCGAAAAGATCAAGAGCAGCCCGCATGGATTTGAC<br>TTGGTCAGGGCCGAGCCTACATGTGCGAATGATGCCCATACTTGAGCCACCTAACTTTGTTTTAGGGC<br>GACTGCCCTGCTGCGTAACATCGTTGCTGCTGCGTAACATCGTTGCTGCTCCATAACATCAAACATCGA<br>CCCACGGCGTAACGCGCTTGCTGCTTGGATGCCCGAGGCATAGACTGTACAAAAAAACAGTCATAACA<br>AGCCATGAAAACCGCCACTGCGCCGTTACCACCGCTGCGTTCGGTCAAGGTTCTGGACCAGTTGCGTG<br>AGCGCATACGCTACTTGCATTACAGTTTACGAACCGAACAGGCTTATGTCAACTGGGTTCGTGCCTTCA<br>TCCGTTTCCACGGTGTGCGTCACCCGGCAACCTTGGGCAGCAGCGAAGTCGCCATAACTTCGTATAGC<br>ATACATTATACGAAGTTATCTGCCAGGCACATGGGTTTTACTAGTATCGATTCGCGACCTACTCCGGAA<br>TATTAATAGATCATGGAGATAATTAAAATGATAACCATCTCGCAAATAAATAAGTATTTTACTGTTTTCG<br>TAACAGTTTTGTAATAAAAAAACCTATAAATATTCCGGATTATTCATACCGTCCCACCATCGGGCGCGG<br>ATCCCGACCATGCATCACCATCACCATCACCATAATCAGTGCGCGAAGGACGCGCGGATCCCGACCAT<br>GCATCACCATCACCATCACCATAATCAGTGCGCGAAGGACATAACTCATGAAGCCTCCAGTATACCCAT<br>CGATTTGCAAGAAAGATACTCGCACTGGAAGAAAAACACTAAACTACTTTATGATTACCTAAACACGA<br>ATTCAACAAAGTGGCCGTCCTTAACGTGCCAGTTCTTTCCTGATTTAGATACCACTTCGGATGAGCATC<br>GCATCTTGTTATCCTCATTTACATCTTCCCAAAAACCTGAAGATGAGACCATATATATTAGCAAAATATC<br>CACGTTGGGTCATATAAAATGGTCATCTTTAAATAATTTCGACATGGACGAAATGGAATTCAAACCGG<br>AGAACTCGACAAGGTTTCCCTCCAAACACTTAGTAAATGACATCAGTATTTTCTTCCCAAACGGGGAAT<br>GCAATAGGGCAAGATATTTGCCTCAAAATCCAGATATTATAGCCGGCGCCTCTTCAGATGGTGCAATCT<br>ACATATTCGATAGAACAAAACACGGCTCTACTAGAATAAGACAGTCCAAAATTTCACATCCCTTTGAGA<br>CAAAGCTGTTTGGTTCACATGGTGTTATTCAAGACGTGGAGGCAATGGATACTTCTTCGGCAGATATA<br>AATGAGGCGACTTCTTTAGCCTGGAACTTGCAGCAGGAGGCCCTTTTACTTTCTTCTCACTCCAACGGC<br>CAAGTTCAAGTTTGGGACATTAAACAATATTCGCATGAGAACCCTATAATAGATTTACCCTTAGTGTCA<br>ATAAACAGCGACGGAACAGCGGTGAATGATGTAACTTGGATGCCAACACACGATTCCCTCTTTGCTGC<br>TTGTACTGAAGGAAATGCGGTCTCCCTATTAGATCTGAGGACTAAGAAAGAGAAGCTCCAGAGTAACC<br>GTGAAAAACACGATGGTGGAGTAAACTCCTGTAGATTTAACTATAAGAACTCTTTAATTCTAGCATCTG<br>CAGATTCAAATGGGAGGCTAAATTTATGGGATATTAGAAACATGAACAAAAGCCCAATCGCTACCATG<br>GAGCACGGTACTTCCGTTTCAACTTTAGAATGGAGTCCAAATTTCGATACTGTATTGGCAACGGCTGGC<br>CAAGAAGATGGGTTAGTCAAGCTATGGGATACCTCCTGCGAAGAAACTATATTTACCCATGGTGGTCA<br>TATGCTCGGTGTGAACGACATTTCGTGGGACGCTCATGACCCTTGGTTAATGTGCAGTGTGGCAAATG<br>ATAATTCAGTTCACATATGGAAACCTGCAGGAAACCTTGTTGGACATTCGTGAGCTCTAGAGCCTGCA<br>GTCTCGACAAGCTTGTCGAGAAGTACTAGAGGATCATAATCAGCCATACCACATTTGTAGAGGTTTTAC<br>TTGCTTTAAAAAACCTCCCACACCTCCCCCTGAACCTGAAACATAAAATGAATGCAATTGTTGTTGTTAA<br>CTTGTTTATTGCAGCTTATAATGGTTACAAATAAAGCAATAGCATCACAAATTTCACAAATAAAGCATTT<br>TTTTCACTGCATTCTAGTTGTGGTTTGTCCAAACTCATCAATGTATCTTATCATGTCTGGATCTGATCACT<br>GCTTGAGCCTAGAAGATCCGGCTGCTAACAAAGCCCGAAAGGAAGCTGAGTTGGCTGCTGCCACCGC<br>TGAGCAATAACTATCATAACCCCTAGGTGCCATTTCATTACCTCTTTCTCCGCACCCGACATAGATCTGG<br>GCCAACTTTTGGCGAAAATGAGACGTTGATCGGCACGTAAGAGGGTCCAACTTTCACCATAATGAAAT<br>AAGATCACTACCGGGCGTATTTTTTGAGTTATCGAGATTTTCAGGAGCTAAGGAAGCTAAAATGGAGA |

| | |
|---|---|
| | AAAAAATCACTGGATATACCACCGTTGATATATCCCAATGGCATCGTAAAGAACATTTTGAGGCATTTC<br>AGTCAGTTGCTCAATGTACCTATAACCAGACCGTTCAGCTGGATATTACGGCCTTTTTAAAGACCGTAA<br>AGAAAAATAAGCACAAGTTTTATCCGGCCTTTATTCACATTCTTGCCCGCCTGATGAATGCTCATCCGG<br>AATTCCGTATGGCAATGAAAGACGGTGAGCTGGTGATATGGGATAGTGTTCACCCTTGTTACACCGTT<br>TTCCATGAGCAAACTGAAACGTTTTCATCGCTCTGGAGTGAATACCACGACGATTTCCGGCAGTTTCTA<br>CACATATATTCGCAAGATGTGGCGTGTTACGGTGAAAACCTGGCCTATTTCCCTAAAGGGTTTATTGAG<br>AATATGTTTTTCGTCTCAGCCAATCCCTGGGTGAGTTTCACCAGTTTTGATTTAAACGTGGCCAATATG<br>GACAACTTCTTCGCCCCCGTTTTCACCATGGGCAAATATTATACGCAAGGCGACAAGGTGCTGATGCC<br>GCTGGCGATTCAGGTTCATCATGCCGTTTGTGATGGCTTCCATGTCGGCAGAATGCTTAATGAATTACA<br>ACAGTACTGCGATGAGTGGCAGGGCGGGGCGTAATTTTTTTAAGGCAGTTATTGGTGCCCTTAAACGC<br>CTGGTTGCTACGCCTGAATAAGTGATAATAAGCGGATGAATGGCAGAAATTCGAAAGCAAATTCGACC<br>CGGTCGTCGGTTCAGGGCAGGGTCGTTAAATAGCCGCTTATGTCTATTGCTGGTTTACCGGTTTATTGA<br>CTACCGGAAGCAGTGTGACCGTGTGCTTCTCAAATGCCTGAGGCCAGTTTGCTCAGGCTCTCCCCGTG<br>GAGGTAATAATTGACGATATGATCATTTATTCTGCCTCCCAGCTGACATTCATCCGGGGTCAGCACCGT<br>TTCTGCGGACTGGCTTTCTACGTGTTCCGCTTCCTTTAGCAGCCCTTGCGCCCTGAGTGCTTGCGGCAG<br>CGTGAAGCTAATTCCCATGTCAGCCGTTAAGTGTTCCTGTGTCACTCAAAATTGCTTTGAGAGGCTCTA<br>AGGGCTTCTCAGTGCGTTACATCCCTGGCTTGTTGTCCACAACCGTTAAACCTTAAAAGCTTTAAAAGC<br>CTTATATATTCTTTTTTTTCTTATAAAACTTAAAACCTTAGAGGCTATTTAAGTTGCTGATTTATATTAATT<br>TTATTGTTCAAACATGAGAGCTTAGTACGTGAAACATGAGAGCTTAGTACGTTAGCCATGAGAGCTTA<br>GTACGTTAGCCATGAGGGTTTAGTTCGTTAAACATGAGAGCTTAGTACGTTAAACATGAGAGCTTAGT<br>ACGTGAAACATGAGAGCTTAGTACGTACTATCAACAGGTTGAACTGCTGATCAACAGATCCTCTACGC<br>GGCCGCGGTACCATAACTTCGTATAGCATACATTATACGAAGTTATCTGTAACTATAACGGTCCTAAGG<br>TAGCGAGTTTAAACACTAGTATCGATTCGCGACCTACTCCGGAATATTAATAGATCATGGAGATAATTA<br>AAATGATAACCATCTCGCAAATAAATAAGTATTTTACTGTTTTCGTAACAGTTTTGTAATAAAAAAACCT<br>ATAAATATTCCGGATTATTCATACCGTCCCACCATCGGGCGCGGATCCCGGTCCGAAGCGCGCGGAAT<br>TCAAAGGCCTACGTCGACGAGCTCACTTGTCGCGGCCGCTTTCGAATCTAGAGCCTGCAGTCTCGACA<br>AGCTTGTCGAGAAGTACTAGAGGATCATAATCAGCCATACCACATTTGTAGAGGTTTTACTTGCTTTAA<br>AAAACCTCCCACACCTCCCCCTGAACCTGAAACATAAAATGAATGCAATTGTTGTTGTTAACTTGTTTAT<br>TGCAGCTTATAATGGTTACAAATAAAGCAATAGCATCACAAATTTCACAAATAAAGCATTTTTTTCACT<br>GCATTCTAGTTGTGGTTTGTCCAAACTCATCAATGTATCTTATCATGTCTGGATCTGATCACTGCTTGAG<br>CCTAGAAGATCCGGCTGCTAACAAAGCCCGAAAGGAAGCTGAGTTGGCTGCTGCCACCGCTGAGCAA<br>TAACTATCATAACCCCTAGGGTATACCCATCTAATTGGAACCAGATAAGTGAAATCTAGTTCCAAACTA<br>TTTTGTCATTTTTAATTTTCGTATTAGCTTACGACGCTACACCCAGTTCCCATCTATTTTGTCACTCTTCCC<br>TAAATAATCCTTAAAAACTCCATTTCCACCCCTCCCAGTTCCCAACTATTTTGTCCGCCCACAACCGGTT<br>GACTTGGGTCAACTGTCAGACCAAGTTTACTCATATATACTTTAGATTGATTTAAAACTTCATTTTTAAT<br>TTAAAAGGATCTAGGTGAAGATCCTTTTTGATAATCTCATGACCAAAATCCCTTAACGTGAGTTTTCGTT<br>CCACTGAGCGTCAGACCCCGTAGAAAAGATCAAAGGATCTTCTTGAGATCCTTTTTTTCTGCGCGTAAT<br>CTGCTGCTTGCAAACAAAAAAACCACCGCTACCAGCGGTGGTTTGTTTGCCGGATCAAGAGCTACCAA<br>CTCTTTTTCCGAAGGTAACTGGCTTCAGCAGAGCGCAGATACCAAATACTGTTCTTCTAGTGTAGCCGT<br>AGTTAGGCCACCACTTCAAGAACTCTGTAGCACCGCCTACATACCTCGCTCTGCTAATCCTGTTACCAGT<br>GGCTGCTGCCAGTGGCGATAAGTCGTGTCTTACCGGGTTGGACTCAAGACGATAGTTACCGGATAAG<br>GCGCAGCGGTCGGGCTGAACGGGGGGTTCGTGCACACAGCCCAGCTTGGAGCG |
| pRS415 | CGCTCCAAGCTGGGCTGTGTGCACGAACCCCCCGTTCAGCCCGACCGCTGCGCCTTATCCGGTAACTAT<br>CGTCTTGAGTCCAACCCGGTAAGACACGACTTATCGCCACTGGCAGCAGCCACTGGTAACAGGATTAG<br>CAGAGCGAGGTATGTAGGCGGTGCTACAGAGTTCTTGAAGTGGTGGCCTAACTACGGCTACACTAGA<br>AGGACAGTATTTGGTATCTGCGCTCTGCTGAAGCCAGTTACCTTCGGAAAAAGAGTTGGTAGCTCTTG<br>ATCCGGCAAACAAACCACCGCTGGTAGCGGTGGTTTTTTTGTTTGCAAGCAGCAGATTACGCGCAGAA<br>AAAAAGGATCTCAAGAAGATCCTTTGATCTTTTCTACGGGGTCTGACGCTCAGTGGAACGAAAACTCA<br>CGTTAAGGGATTTTGGTCATGAGATTATCAAAAAGGATCTTCACCTAGATCCTTTTAAATTAAAAATGA<br>AGTTTTAAATCAATCTAAAGTATATATGAGTAAACTTGGTCTGACAGTTACCAATGCTTAATCAGTGAG<br>GCACCTATCTCAGCGATCTGTCTATTTCGTTCATCCATAGTTGCCTGACTCCCCGTCGTGTAGATAACTA<br>CGATACGGGAGGGCTTACCATCTGGCCCCAGTGCTGCAATGATACCGCGAGACCCACGCTCACCGGCT<br>CCAGATTTATCAGCAATAAACCAGCCAGCCGGAAGGGCCGAGCGCAGAAGTGGTCCTGCAACTTTATC<br>CGCCTCCATCCAGTCTATTAATTGTTGCCGGGAAGCTAGAGTAAGTAGTTCGCCAGTTAATAGTTTGCG<br>CAACGTTGTTGCCATTGCTACAGGCATCGTGGTGTCACGCTCGTCGTTTGGTATGGCTTCATTCAGCTC |

| | |
|---|---|
| | CGGTTCCCAACGATCAAGGCGAGTTACATGATCCCCCATGTTGTGCAAAAAAGCGGTTAGCTCCTTCG<br>GTCCTCCGATCGTTGTCAGAAGTAAGTTGGCCGCAGTGTTATCACTCATGGTTATGGCAGCACTGCATA<br>ATTCTCTTACTGTCATGCCATCCGTAAGATGCTTTTCTGTGACTGGTGAGTACTCAACCAAGTCATTCTG<br>AGAATAGTGTATGCGGCGACCGAGTTGCTCTTGCCCGGCGTCAATACGGGATAATACCGCGCCACATA<br>GCAGAACTTTAAAAGTGCTCATCATTGGAAAACGTTCTTCGGGGCGAAAACTCTCAAGGATCTTACCG<br>CTGTTGAGATCCAGTTCGATGTAACCCACTCGTGCACCCAACTGATCTTCAGCATCTTTTACTTTCACCA<br>GCGTTTCTGGGTGAGCAAAAACAGGAAGGCAAAATGCCGCAAAAAAGGGAATAAGGGCGACACGGA<br>AATGTTGAATACTCATACTCTTCCTTTTTCAATATTATTGAAGCATTTATCAGGGTTATTGTCTCATGAGC<br>GGATACATATTTGAATGTATTTAGAAAAATAAACAAATAGGGGTTCCGCGCACATTTCCCCGAAAAGT<br>GCCACCTGGGTCCTTTTCATCACGTGCTATAAAAATAATTATAATTTAAATTTTTTAATATAAATATATA<br>AATTAAAAATAGAAAGTAAAAAAAGAAATTAAAGAAAAAATAGTTTTTGTTTTCCGAAGATGTAAAAG<br>ACTCTAGGGGGATCGCCAACAAATACTACCTTTTATCTTGCTCTTCCTGCTCTCAGGTATTAATGCCGAA<br>TTGTTTCATCTTGTCTGTGTAGAAGACCACACACGAAAATCCTGTGATTTTACATTTTACTTATCGTTAA<br>TCGAATGTATATCTATTTAATCTGCTTTTCTTGTCTAATAAATATATATGTAAAGTACGCTTTTTGTTGAA<br>ATTTTTTAAACCTTTGTTTATTTTTTTTTCTTCATTCCGTAACTCTTCTACCTTCTTTATTTACTTTCTAAAA<br>TCCAAATACAAAACATAAAAATAAATAAACACAGAGTAAATTCCCAAATTATTCCATCATTAAAAGATA<br>CGAGGCGCGTGTAAGTTACAGGCAAGCGATCCGTCCTAAGAAACCATTATTATCATGACATTAACCTA<br>TAAAAATAGGCGTATCACGAGGCCCTTTCGTCTCGCGCGTTTCGGTGATGACGGTGAAAACCTCTGAC<br>ACATGCAGCTCCCGGAGACGGTCACAGCTTGTCTGTAAGCGGATGCCGGGAGCAGACAAGCCCGTCA<br>GGGCGCGTCAGCGGGTGTTGGCGGGTGTCGGGGCTGGCTTAACTATGCGGCATCAGAGCAGATTGTA<br>CTGAGAGTGCACCATATCGACTACGTCGTTAAGGCCGTTTCTGACAGAGTAAAATTCTTGAGGGAACT<br>TTCACCATTATGGGAAATGGTTCAAGAAGGTATTGACTTAAACTCCATCAAATGGTCAGGTCATTGAGT<br>GTTTTTTATTTGTTGTATTTTTTTTTTTTTAGAGAAAATCCTCCAATATATAAATTAGGAATCATAGTTTC<br>ATGATTTTCTGTTACACCTAACTTTTTGTGTGGTGCCCTCCTCCTTGTCAATATTAATGTTAAAGTGCAAT<br>TCTTTTTCCTTATCACGTTGAGCCATTAGTATCAATTTGCTTACCTGTATTCCTTTACATCCTCCTTTTTCT<br>CCTTCTTGATAAATGTATGTAGATTGCGTATATAGTTTCGTCTACCCTATGAACATATTCCATTTTGTAAT<br>TTCGTGTCGTTTCTATTATGAATTTCATTTATAAAGTTTATGTACAAATATCATAAAAAAAGAGAATCTT<br>TTTAAGCAAGGATTTTCTTAACTTCTTCGGCGACAGCATCACCGACTTCGGTGGTACTGTTGGAACCAC<br>CTAAATCACCAGTTCTGATACCTGCATCCAAAACCTTTTTAACTGCATCTTCAATGGCCTTACCTTCTTCA<br>GGCAAGTTCAATGACAATTTCAACATCATTGCAGCAGACAAGATAGTGGCGATAGGGTTGACCTTATT<br>CTTTGGCAAATCTGGAGCAGAACCGTGGCATGGTTCGTACAAACCAAATGCGGTGTTCTTGTCTGGCA<br>AAGAGGCCAAGGACGCAGATGGCAACAAACCCAAGGAACCTGGGATAACGGAGGCTTCATCGGAGA<br>TGATATCACCAAACATGTTGCTGGTGATTATAATACCATTTAGGTGGGTTGGGTTCTTAACTAGGATCA<br>TGGCGGCAGAATCAATCAATTGATGTTGAACCTTCAATGTAGGGAATTCGTTCTTGATGGTTTCCTCCA<br>CAGTTTTTCTCCATAATCTTGAAGAGGCCAAAACATTAGCTTTATCCAAGGACCAAATAGGCAATGGTG<br>GCTCATGTTGTAGGGCCATGAAAGCGGCCATTCTTGTGATTCTTTGCACTTCTGGAACGGTGTATTGTT<br>CACTATCCCAAGCGACACCATCACCATCGTCTTCCTTTCTCTTACCAAAGTAAATACCTCCCACTAATTCT<br>CTGACAACAACGAAGTCAGTACCTTTAGCAAATTGTGGCTTGATTGGAGATAAGTCTAAAAGAGAGTC<br>GGATGCAAAGTTACATGGTCTTAAGTTGGCGTACAATTGAAGTTCTTTACGGATTTTTAGTAAACCTTG<br>TTCAGGTCTAACACTACCGGTACCCCATTTAGGACCACCCACAGCACCTAACAAAACGGCATCAGCCTT<br>CTTGGAGGCTTCCAGCGCCTCATCTGGAAGTGGAACACCTGTAGCATCGATAGCAGCACCACCAATTA<br>AATGATTTTCGAAATCGAACTTGACATTGGAACGAACATCAGAAATAGCTTTAAGAACCTTAATGGCTT<br>CGGCTGTGATTTCTTGACCAACGTGGTCACCTGGCAAAACGACGATCTTCTTAGGGGCAGACATAGGG<br>GCAGACATTAGAATGGTATATCCTTGAAATATATATATATATATTGCTGAAATGTAAAAGGTAAGAAA<br>AGTTAGAAAGTAAGACGATTGCTAACCACCTATTGGAAAAAACAATAGGTCCTTAAATAATATTGTCA<br>ACTTCAAGTATTGTGATGCAAGCATTTAGTCATGAACGCTTCTCTATTCTATATGAAAAGCCGGTTCCG<br>GCGCTCTCACCTTTCCTTTTTCTCCCAATTTTTCAGTTGAAAAAGGTATATGCGTCAGGCGACCTCTGAA<br>ATTAACAAAAAATTTCCAGTCATCGAATTTGATTCTGTGCGATAGCGCCCCTGTGTGTTCTCGTTATGTT<br>GAGGAAAAAAATAATGGTTGCTAAGAGATTCGAACTCTTGCATCTTACGATACCTGAGTATTCCCACA<br>GTTAACTGCGGTCAAGATATTTCTTGAATCAGGCGCCTTAGACCGCTCGGCCAAACAACCAATTACTTG<br>TTGAGAAATAGAGTATAATTATCCTATAAATATAACGTTTTTGAACACACATGAACAAGGAAGTACAG<br>GACAATTGATTTTGAAGAGAATGTGGATTTTGATGTAATTGTTGGGATTCCATTTTTAATAAGGCAATA<br>ATATTAGGTATATGGATATACTAGAAGTTCTCCTCGACCGGTCGATATGCGGTGTGAAATACCGCACA<br>GATGCGTAAGGAGAAAATACCGCATCAGGAAATTGTAAGCGTTAATATTTTGTTAAAATTCGCGTTAA<br>ATTTTTGTTAAATCAGCTCATTTTTTAACCAATAGGCCGAAATCGGCAAAATCCCTTATAAATCAAAAGA<br>ATAGACCGAGATAGGGTTGAGTGTTGTTCCAGTTTGGAACAAGAGTCCACTATTAAAGAACGTGGACT<br>CCAACGTCAAAGGGCGAAAAACCGTCTATCAGGGCGATGGCCCACTACGTGAACCATCACCCTAATCA |

| | |
|---|---|
| | AGTTTTTTGGGGTCGAGGTGCCGTAAAGCACTAAATCGGAACCCTAAAGGGAGCCCCCGATTTAGAGC<br>TTGACGGGGAAAGCCGGCGAACGTGGCGAGAAAGGAAGGGAAGAAAGCGAAAGGAGCGGGCGCTA<br>GGGCGCTGGCAAGTGTAGCGGTCACGCTGCGCGTAACCACCACACCCGCCGCGCTTAATGCGCCGCT<br>ACAGGGCGCGTCCATTCGCCATTCAGGCTGCGCAACTGTTGGGAAGGGCGATCGGTGCGGGCCTCTT<br>CGCTATTACGCCAGCTGGCGAAAGGGGGATGTGCTGCAAGGCGATTAAGTTGGGTAACGCCAGGGTT<br>TTCCCAGTCACGACGTTGTAAAACGACGGCCAGTGAGCGCGCGTAATACGACTCACTATAGGGCGAAT<br>TGGGTACCGGGCCCCCCCTCGAGGTCGACGGTATCGATAAGCTTGATATCGAATTCCTGCAGCCCGGG<br>GGATCCACTAGTTCTAGAGCGGCCGCCACCGCGGTGGAGCTCCAGCTTTTGTTCCCTTTAGTGAGGGT<br>TAATTGCGCGCTTGGCGTAATCATGGTCATAGCTGTTTCCTGTGTGAAATTGTTATCCGCTCACAATTCC<br>ACACAACATACGAGCCGGAAGCATAAAGTGTAAAGCCTGGGGTGCCTAATGAGTGAGCTAACTCACA<br>TTAATTGCGTTGCGCTCACTGCCCGCTTTCCAGTCGGGAAACCTGTCGTGCCAGCTGCATTAATGAATC<br>GGCCAACGCGCGGGGAGAGGCGGTTTGCGTATTGGGCGCTCTTCCGCTTCCTCGCTCACTGACTCGCT<br>GCGCTCGGTCGTTCGGCTGCGGCGAGCGGTATCAGCTCACTCAAAGGCGGTAATACGGTTATCCACAG<br>AATCAGGGGATAACGCAGGAAAGAACATGTGAGCAAAAGGCCAGCAAAAGGCCAGGAACCGTAAAA<br>AGGCCGCGTTGCTGGCGTTTTTCCATAGGCTCCGCCCCCCTGACGAGCATCACAAAAATCGACGCTCA<br>AGTCAGAGGTGGCGAAACCCGACAGGACTATAAAGATACCAGGCGTTTCCCCCTGGAAGCTCCCTCGT<br>GCGCTCTCCTGTTCCGACCCTGCCGCTTACCGGATACCTGTCCGCCTTTCTCCCTTCGGGAAGCGTGGC<br>GCTTTCTCATAGCTCACGCTGTAGGTATCTCAGTTCGGTGTAGGTCGTT |
| 207bp loading control | ATCTAGTATTAATTAATATGAATTCGGATCCACATGCACAGGATGTATATATCTGACACGTGCCTGGAG<br>ACTAGGGAGTAATCCCCTTGGCGGTTAAAACGCGGGGGACAGCGCGTACGTGCGTTTAAGCGGTGCT<br>AGAGCTGTCTACGACCAATTGAGCGGCCTCGGCACCGGGATTCTCCAGGGCGGCCGCGTATAGGGTC<br>CGAT |

Supplementary Table S3: - adapters used during MNase-seq (PCNA-NAQ assay)

| | Sequence |
|---|---|
| A1_AACACCTA_FWD | ACACTCTTTCCCTACACGACGCTCTTCCGATCTAACACCTA*T |
| A1_AACACCTA_REV | /5Phos/TAGGTGTTAGATCGGAAGAGCGGTTCAGCAGGAATGCCGAG |
| A2_ACGTAGCT_FWD | ACACTCTTTCCCTACACGACGCTCTTCCGATCTACGTAGCT*T |
| A2_ACGTAGCT_REV | /5Phos/AGCTACGTAGATCGGAAGAGCGGTTCAGCAGGAATGCCGAG |
| A3_ATATAGGA_FWD | ACACTCTTTCCCTACACGACGCTCTTCCGATCTATATAGGA*T |
| A3_ATATAGGA_REV | /5Phos/TCCTATATAGATCGGAAGAGCGGTTCAGCAGGAATGCCGAG |
| A4_CACAGTTG_FWD | ACACTCTTTCCCTACACGACGCTCTTCCGATCTCACAGTTG*T |
| A4_CACAGTTG_REV | /5Phos/CAACTGTGAGATCGGAAGAGCGGTTCAGCAGGAATGCCGAG |
| A5_CCTACAAC_FWD | ACACTCTTTCCCTACACGACGCTCTTCCGATCTCCTACAAC*T |
| A5_CCTACAAC_REV | /5Phos/GTTGTAGGAGATCGGAAGAGCGGTTCAGCAGGAATGCCGAG |
| A6_CGTCGGCT_FWD | ACACTCTTTCCCTACACGACGCTCTTCCGATCTCGTCGGCT*T |
| A6_CGTCGGCT_REV | /5Phos/AGCCGACGAGATCGGAAGAGCGGTTCAGCAGGAATGCCGAG |
| A7_GACGTCAA_FWD | ACACTCTTTCCCTACACGACGCTCTTCCGATCTGACGTCAA*T |
| A7_GACGTCAA_REV | /5Phos/TTGACGTCAGATCGGAAGAGCGGTTCAGCAGGAATGCCGAG |
| A8_GCGTTTCG_FWD | ACACTCTTTCCCTACACGACGCTCTTCCGATCTGCGTTTCG*T |
| A8_GCGTTTCG_REV | /5Phos/CGAAACGCAGATCGGAAGAGCGGTTCAGCAGGAATGCCGAG |

Supplementary Table S4, DNA sequences of the plasmid used for primer extension experiments:

| Name | DNA sequence |
|---|---|
| pBluescript SK(-)-pC3N sequence: | CACCTGACGCGCCCTGTAGCGGCGCATTAAGCGCGGCGGGTGTGGTGGTTACGCGCAGCGTGACCG<br>CTACACTTGCCAGCGCCCTAGCGCCCGCTCCTTTCGCTTTCTTCCCTTCCTTTCTCGCCACGTTCGCCG<br>GCTTTCCCCGTCAAGCTCTAAATCGGGGGCTCCCTTTAGGGTTCCGATTTAGTGCTTTACGGCACCTC<br>GACCCCAAAAAACTTGATTAGGGTGATGGTTCACGTAGTGGGCCATCGCCCTGATAGACGGTTTTTC<br>GCCCTTTGACGTTGGAGTCCACGTTCTTTAATAGTGGACTCTTGTTCCAAACTGGAACAACACTCAAC<br>CCTATCTCGGTCTATTCTTTTGATTTATAAGGGATTTTGCCGATTTCGGCCTATTGGTTAAAAAATGAG<br>CTGATTTAACAAAAATTTAACGCGAATTTTAACAAAATATTAACGCTTACAATTTCCATTCGCCATTCA<br>GGCTGCGCAACTGTTGGGAAGGGCGATCGGTGCGGGCCTCTTCGCTATTACGCCAGCTGGCGAAAG<br>GGGGATGTGCTGCAAGGCGATTAAGTTGGGTAACGCCAGGGTTTTCCCAGTCACGACGTTGTAAAA<br>CGACGGCCAGTGAATTGTAATACGACTCACTATAGGGCGAATTGGGTACCGGGCCCCCCCTCGAGG<br>TCGACGGTATCGATAaGCTTGGGAcccTGGGAGGGAGATCCACTAGTTCTAGAGCGGCCGCCACCGC<br>GGTGGAGCTCCAGCTTTTGTTCCCTTTAGTGAGGGTTAATTTCGAGCTTGGCGTAATCATGGTCATA<br>GCTGTTTCCTGTGTGAAATTGTTATCCGCTCACAATTCCACACAACATACGAGCCGGAAGCATAAAGT<br>GTAAAGCCTGGGGTGCCTAATGAGTGAGCTAACTCACATTAATTGCGTTGCGCTCACTGCCCGCTTT<br>CCAGTCGGGAAACCTGTCGTGCCAGCTGCATTAATGAATCGGCCAACGCGCGGGGAGAGGCGGTTT<br>GCGTATTGGGCGCTCTTCCGCTTCCTCGCTCACTGACTCGCTGCGCTCGGTCGTTCGGCTGCGGCGA<br>GCGGTATCAGCTCACTCAAAGGCGGTAATACGGTTATCCACAGAATCAGGGGATAACGCAGGAAAG<br>AACATGTGAGCAAAAGGCCAGCAAAAGGCCAGGAACCGTAAAAAGGCCGCGTTGCTGGCGTTTTTC<br>CATAGGCTCCGCCCCCCTGACGAGCATCACAAAAATCGACGCTCAAGTCAGAGGTGGCGAAACCCG<br>ACAGGACTATAAAGATACCAGGCGTTTCCCCCTGGAAGCTCCCTCGTGCGCTCTCCTGTTCCGACCCT<br>GCCGCTTACCGGATACCTGTCCGCCTTTCTCCCTTCGGGAAGCGTGGCGCTTTCTCATAGCTCACGCT<br>GTAGGTATCTCAGTTCGGTGTAGGTCGTTCGCTCCAAGCTGGGCTGTGTGCACGAACCCCCCGTTCA<br>GCCCGACCGCTGCGCCTTATCCGGTAACTATCGTCTTGAGTCCAACCCGGTAAGACACGACTTATCG<br>CCACTGGCAGCAGCCACTGGTAACAGGATTAGCAGAGCGAGGTATGTAGGCGGTGCTACAGAGTTC<br>TTGAAGTGGTGGCCTAACTACGGCTACACTAGAAGGACAGTATTTGGTATCTGCGCTCTGCTGAAGC<br>CAGTTACCTTCGGAAAAAGAGTTGGTAGCTCTTGATCCGGCAAACAAACCACCGCTGGTAGCGGTG<br>GTTTTTTTGTTTGCAAGCAGCAGATTACGCGCAGAAAAAAAGGATCTCAAGAAGATCCTTTGATCTTT<br>TCTACGGGGTCTGACGCTCAGTGGAACGAAAACTCACGTTAAGGGATTTTGGTCATGAGATTATCAA<br>AAAGGATCTTCACCTAGATCCTTTTAAATTAAAAATGAAGTTTTAAATCAATCTAAAGTATATATGAG<br>TAAACTTGGTCTGACAGTTACCAATGCTTAATCAGTGAGGCACCTATCTCAGCGATCTGTCTATTTCG<br>TTCATCCATAGTTGCCTGACTCCCCGTCGTGTAGATAACTACGATACGGGAGGGCTTACCATCTGGCC<br>CCAGTGCTGCAATGATACCGCGAGACCCACGCTCACCGGCTCCAGATTTATCAGCAATAAACCAGCC<br>AGCCGGAAGGGCCGAGCGCAGAAGTGGTCCTGCAACTTTATCCGCCTCCATCCAGTCTATTAATTGT<br>TGCCGGGAAGCTAGAGTAAGTAGTTCGCCAGTTAATAGTTTGCGCAACGTTGTTGCCATTGCTACAG<br>GCATCGTGGTGTCACGCTCGTCGTTTGGTATGGCTTCATTCAGCTCCGGTTCCCAACGATCAAGGCG<br>AGTTACATGATCCCCCATGTTGTGCAAAAAAGCGGTTAGCTCCTTCGGTCCTCCGATCGTTGTCAGAA |

| | |
|---|---|
| | GTAAGTTGGCCGCAGTGTTATCACTCATGGTTATGGCAGCACTGCATAATTCTCTTACTGTCATGCCA<br>TCCGTAAGATGCTTTTCTGTGACTGGTGAGTACTCAACCAAGTCATTCTGAGAATAGTGTATGCGGC<br>GACCGAGTTGCTCTTGCCCGGCGTCAATACGGGATAATACCGCGCCACATAGCAGAACTTTAAAAGT<br>GCTCATCATTGGAAAACGTTCTTCGGGGCGAAAACTCTCAAGGATCTTACCGCTGTTGAGATCCAGT<br>TCGATGTAACCCACTCGTGCACCCAACTGATCTTCAGCATCTTTTACTTTCACCAGCGTTTCTGGGTGA<br>GCAAAAACAGGAAGGCAAAATGCCGCAAAAAAGGGAATAAGGGCGACACGGAAATGTTGAATACT<br>CATACTCTTCCTTTTTCAATATTATTGAAGCATTTATCAGGGTTATTGTCTCATGAGCGGATACATATT<br>TGAATGTATTTAGAAAAATAAACAAATAGGGGTTCCGCGCACATTTCCCCGAAAAGTGC |

Supplementary Table S5: Primers used for establishing the *Chaf1a*-dTAG cell line.

| | Sequence |
|---|---|
| sgRNA#1 | CGCCGTCGCGGAGATGTTGG *AGG* |
| Primer#1 | CAATGGCTACTTTCAACCCGTC |
| Primer#2 | CACCCAAACCGACCTTCCTG |
| Primer#3 | GACGTACTGAGTGCACCTCTT |
| Primer#4 | CCAGCCCCTCAATCGTTCAA |
